# Diverse phage defence systems define West African South American pandemic *Vibrio cholerae*

**DOI:** 10.1101/2024.11.23.624991

**Authors:** David W. Adams, Milena Jaskólska, Alexandre Lemopoulos, Sandrine Stutzmann, Laurie Righi, Loriane Bader, Melanie Blokesch

**Author notes:** These authors contributed equally (David W. Adams, Melanie Blokesch).

## Abstract

Our understanding of the factors underlying the evolutionary success of different lineages of pandemic *Vibrio cholerae* remains incomplete. Interestingly, two unique genetic signatures define the West African South American (WASA) lineage of *V. cholerae* responsible for the 1991-2001 Latin American cholera epidemic. Here we show these signatures encode diverse anti-phage defence systems. Firstly, the WASA-1 prophage encodes a 2-gene abortive-infection system WonAB that renders the lineage resistant to the major predatory vibriophage ICP1, which alongside other phages, is thought to restrict cholera epidemics and has potential for use in prophylaxis. Secondly, a unique set of genes on the *Vibrio* seventh pandemic island II encodes an unusual modification-dependent restriction system targeting phages with modified genomes, and a new member of the Shedu defence family that defends against vibriophage X29. Taken together, we propose that these anti-phage defence systems have likely contributed to the success of a major epidemic lineage of the ongoing seventh cholera pandemic.

---

The ability of lytic bacteriophages to rapidly devastate bacterial populations has driven the evolution of multiple layers of defence, including a diverse array of specialised anti-phage defence systems^1^. Bacteriophages also have the potential to affect bacterial pathogenesis, as exemplified by the cholera toxin-encoding prophage CTXΦ^2^. Moreover, bacteriophage predation is thought to limit the duration and severity of cholera epidemics, as well as affecting individual patient outcomes^3–6^. Sampling of cholera patients has revealed that the O1 El Tor strains of *Vibrio cholerae* responsible for the ongoing seventh cholera pandemic (7PET) consistently co-occur with three lytic phages ICP1, 2 and 3, with ICP1 being most frequently isolated^7,8^. Furthermore, the demonstration that a cocktail of ICP1-2-3 can prevent cholera in animal infection models has led to renewed interest in using phages as a prophylactic treatment^9^. There is therefore an urgent need to understand the mechanisms by which *V. cholerae* defends against these and other viruses.

Since 1961 7PET strains have spread out from the Bay of Bengal in a series of three distinct but overlapping waves^10^. Importantly, from Wave 2 onwards strains acquired SXT/R391 integrative and conjugative elements (SXT-ICE), which in addition to carrying multiple antibiotic resistance genes, encode a variable anti-phage defence hotspot that in the globally dominant SXT-ICE harbours either BREX or Restriction-Modification systems active against ICP1-3^10,11^. Additionally, a family of viral satellites, the Phage-inducible chromosomal island-like elements (PLE), also occur sporadically and specifically parasitise ICP1 infection to mediate their own transmission^12,13^. Notably, these elements exemplify the co-evolutionary arms race of defence and counter-defence between bacterial hosts and their viral predators, with the emergence of resistant ICP1 phages that can overcome these defence mechanisms selecting for either alternative SXT-ICE carrying new defence systems or new PLE variants^11,14^. In contrast, how 7PET strains that lack SXT-ICE/PLE defend against these phages remains unknown. Given their ubiquitous nature and ability to shape epidemics, we hypothesised that these strains likely contained additional defence systems with activity against ICP1-3.

## WASA-1 prophage renders Peruvian strains resistant to ICP1

To test this hypothesis, we challenged a set of intensively studied wave 1 strains that lack SXT-ICE/PLE with a well-characterised collection of ICP1-3 phages that utilise the O1-antigen (ICP1/3) and OmpU (ICP2) as receptors^7,15,16^. All strains exhibited the expected sensitivities to ICP2 and ICP3 (Fig. 1a and Extended Data Fig. 1a-b). Unexpectedly, however, the widely used model strain A1552 – a Peruvian isolate – was completely resistant to all 16 tested ICP1 isolates spanning a twenty-seven-year period, including the most recent isolates from the Democratic Republic of the Congo^17^, displaying a 10^5^-fold reduction in plaque formation (Fig. 1a and Extended Data Fig. 1a-e). Notably, the contemporary Peruvian strains C6706 and C6709 also behaved similarly. Interestingly, although zones of lysis were still observed at the highest concentrations of phage tested, re-streak tests revealed that these spots contained little-to-no viable particles (Extended Data Fig. 1c), suggesting that ICP1 propagation is significantly impaired and that these lysis zones represent ‘lysis from without’^18^. Indeed, liquid replication assays comparing A1552 with the non-Peruvian strain E7946, showed that although genome injection proceeded similarly in both strains, in contrast to the robust replication observed with E7946, ICP1 was completely unable to replicate on strain A1552 (Extended Data Fig. 1d). ICP1 resistance in strain A1552 was independent of the *Vibrio* pathogenicity islands (VPI-1/2) and the *Vibrio* seventh pandemic islands (VSP-I/II), which together harbour multiple known anti-viral defence systems^19–26^ (Extended Data Fig. 1e). Moreover, the ICP1-resistant strains encode the same repertoire of known defence systems as the ICP1-sensitive strains (Supplementary Table 1), suggesting that a novel as yet unrecognised defence system(s) is likely responsible for ICP1 resistance.

**Fig. 1.**
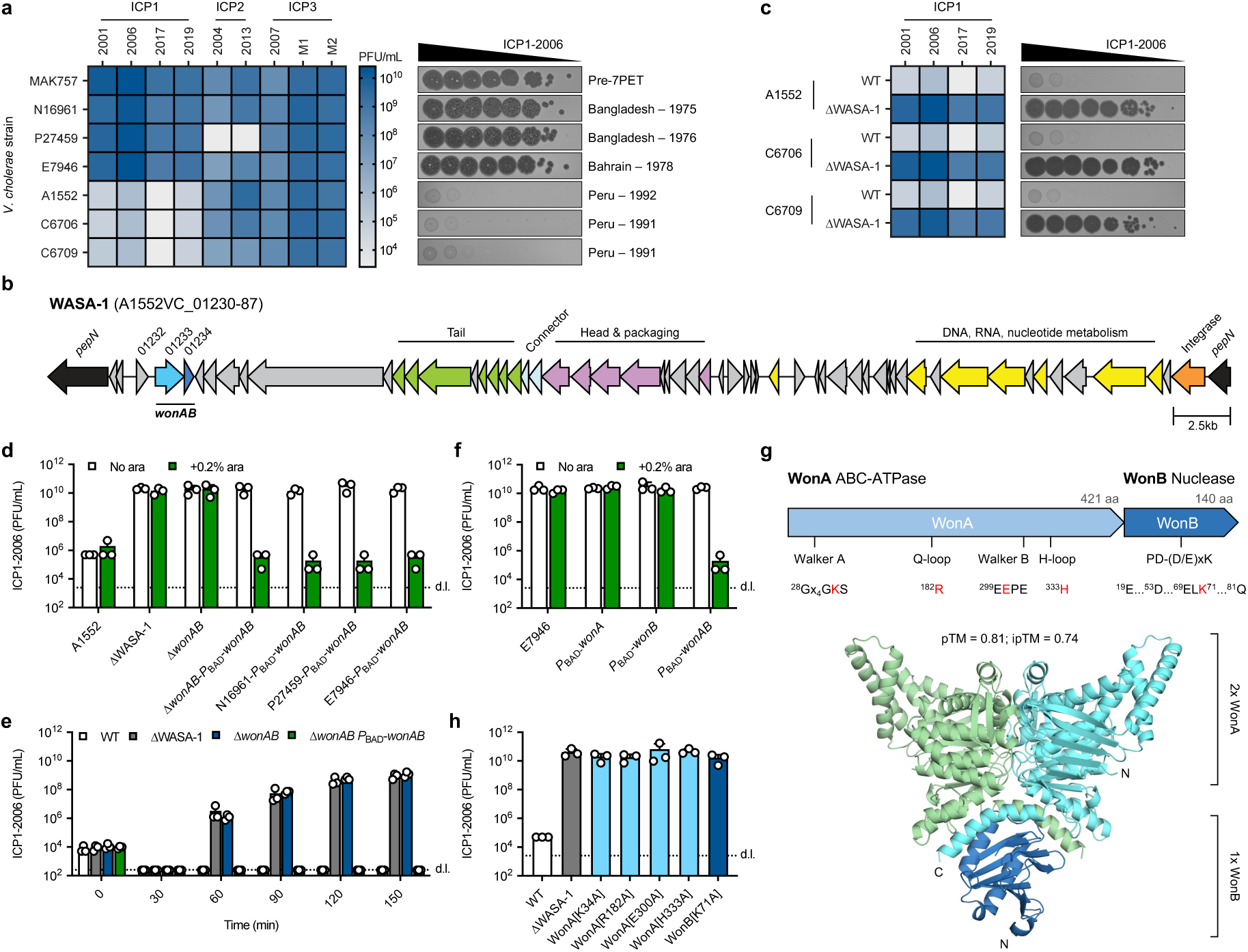
WonAB carried on WASA-1 prophage renders WASA lineage ICP1 resistant. (**a**) Heat-map showing mean plaque-forming units (PFU/mL) of diverse ICP1-3 isolates determined by plaque assays on wave 1 *V. cholerae* strains, as compared to a pre-7^th^ pandemic control (MAK757), shown alongside 10-fold serial dilution plaque assays of ICP1-2006. Data represent the results of three independent repeats. Strain P27459 encodes a naturally occurring resistant variant of the ICP2 receptor OmpU^80^. (**b**) Schematic of the WASA-1 prophage highlighting *wonAB* and genes with predicted functions in prophage biology. (**c**) Heat-map showing mean PFU/mL of diverse ICP1 isolates determined by plaque assays on Peruvian *V. cholerae* strains in WT and ΔWASA-1 backgrounds, determined as in (**a**). (**d**) Plaque assay showing the effects of ΔWASA-1, Δ*wonAB* and *wonAB* complementation in strain A1552 on ICP1-2006 PFU/mL alongside the effect of producing WonAB in non-WASA strains. (**e**) Replication assay showing change in PFU/mL over time following infection of the indicated A1552 cultures with *c.a.* 10^3^-10^4^ PFU of ICP1-2006. (**f**) Plaque assay showing effect of E7946 derivatives producing either WonA, WonB or WonAB on ICP1-2006 PFU/mL. Where indicated, strains in (**d**-**f**) contain a chromosomally integrated transposon carrying the arabinose-inducible *P*_BAD_-promoter, induced by the addition of 0.2% arabinose. (**g**) AlphaFold3 predicted structure of the putative WonAB complex shown below schematic (top) indicating the conserved motifs. Residues targeted by site-directed mutagenesis are highlighted in red. (**h**) Plaque assay testing the effect of WonA and WonB site-directed variants expressed from their native locus on ICP1-2006 PFU/mL. Bar charts show the mean + s.d. from 3 independent repeats. d.l. = detection limit.

The Peruvian strains are members of the West African South American (WASA) lineage of 7PET *V. cholerae*^10^, which was responsible for a massive cholera epidemic that started in Peru in 1991, before spreading rapidly throughout Latin America, resulting in over 1.2 million cholera cases and approximately 12,000 deaths^27^. Interestingly, phylogenetic analysis has shown that these strains originated in West Africa, where they acquired two genetic signatures that define the WASA lineage: (i) a novel set of genes on VSP-II and (ii) the WASA-1 prophage^10,28–31^. Since deletion of VSP-II had no effect on ICP1 resistance we tested strains deleted for WASA-1. Strikingly, this abolished defence against ICP1 in all tested Peruvian strains, and restored ICP1 replication to levels similar to those of a susceptible control (Fig. 1b-e and Extended Data Fig. 1b). Moreover, introducing WASA-1 into non-WASA lineage strains rendered them ICP1 resistant (Extended Data Fig. 1f).

## WonAB system is responsible for ICP1 defence

The WASA-1 prophage is highly conserved throughout all WASA lineage strains^10,30^, and interestingly, is also found in diverse *Vibrio* species (Fig. 1b, Extended Data Fig. 2a and Supplementary Table 2). WASA-1 displays a conserved core genome encoding proteins involved in capsid/tail biogenesis, DNA replication and a recombinase (Fig. 1b). To determine the gene(s) responsible for ICP1 defence, we used an existing RNA-seq dataset^32^ to identify WASA-1 genes expressed under laboratory conditions, and then screened truncations of these regions for loss of protection (Extended Data Fig. 2b). This revealed a 2-gene operon (A1552VC_01233-34) encoding an ATPase and a hypothetical protein, downstream of a conserved putative transcriptional regulator (A1552VC_01232) (Fig. 1b). For the reasons outlined below we have renamed this operon <u>W</u>ASA <u>O</u>LD-ABC ATPase <u>N</u>uclease, WonAB. Strikingly, deletion of *wonAB* abolished protection against ICP1. Moreover, ectopic expression of *wonAB* complemented the Δ*wonAB* deletion and conferred complete protection to ICP1-sensitive non-WASA strains (Fig. 1d-e). Importantly, both genes were required for protection indicating that production of WonAB is both necessary and sufficient for anti-phage activity (Fig. 1f). Notably, although WonAB is unique to the WASA-1 prophage found in *V. cholerae*, known defence systems could also be found at the same locus in other non-cholerae WASA-1 (Supplementary Table 2), consistent with the observation that prophages often carry anti-phage defence systems to increase the fitness of their bacterial hosts^33–35^.

Bioinformatic analysis revealed that WonA is an ATP-binding cassette (ABC) ATPase with a predicted core β-barrel like ATPase fold characteristic of the ABC-ATPase clade^36^. Motifs required for ATP binding and hydrolysis (Walker A, Q-loop, ABC-signature, Walker B, D-loop and H-loop) were readily identifiable (Fig. 1g and Extended Data Fig. 3a). Moreover, structural modelling confidently predicted that WonA forms a dimer, in which these motifs are positioned at the dimer interface to form the composite ATP-binding sites typical of ABC-ATPases^36^ (Fig. 1g, Extended Data Figs. 3b and 4a-b). In contrast, WonB is predicted to encode a variant of the PD-(D/E)xK nuclease fold, with the highly conserved active site residues clustered together^37^ (Fig. 1g and Extended Data Fig. 3c-d). Variants designed to disrupt either WonA ATP-binding/hydrolysis (K34A; R182A; E300A; H333A) or WonB nuclease activity (K71A) all abolished protection against ICP1, despite mostly being produced at similar levels to the WT control (Fig. 1h and Extended Data Fig. 3e). Interestingly, Western blotting showed that inactivating WonA ATPase activity resulted in the loss of WonB, suggesting that WonA regulates or otherwise affects the stability of WonB (Extended Data Fig. 3e). Indeed, the two proteins are predicted to form a complex, with WonB binding to a surface created by the dimerised C-terminal extensions of WonA (Fig. 1g and Extended Data Fig. 4a-d).

Homologues of WonAB are found in diverse classes of mainly Gram-negative bacteria and overall were present in 0.43% of bacterial genomes examined (Extended Data Fig. 3f and Supplementary Table 3). Notably, when expressed in *Escherichia coli* WonAB displayed robust activity against all members of the *Vequintavirinae* subfamily of the BASEL bacteriophage collection^38^, indicating that WonAB activity is not limited to ICP1 (Extended Data Fig. 5). Interestingly, WonA ATPase motifs display deviations typical of the overcoming lysogenisation defect (OLD) family of ABC-ATPases^36,39^ (Extended Data Fig. 3a). However, members of this family such as OLD, Gabija, PARIS and the related Septu, are distinct from WonA in both structure and domain composition^39–44^. In contrast, WonAB appears to be closely related to an operon encoding a predicted OLD-ABC ATPase and a novel version of the restriction-enzyme fold (Extended Data Fig. 4), detected during a recent comprehensive *in silico* survey of ABC-ATPases^36^. Moreover, homologues of this operon are found in 0.96% of bacterial genomes and encompass all detected homologues of WonAB (Supplementary Table 3). Furthermore, WonAB also shows similarity to operons with a similar organisation such as PD-T4-4 that have also recently been identified (Extended Data Fig. 4), albeit with variations in the Insert 1/2 regions and C-terminal extensions of the ATPase, and in some cases a distinct nuclease^34,45^. Taken together, these data suggest that WonAB is part of a larger as yet uncharacterised family of ABC-ATPase sensors that are paired with nuclease effectors.

## ICP1 defence occurs via lysis-independent abortive infection

To gain insights into how WonAB mediates ICP1 defence we monitored the growth kinetics of ICP1-infected cells at different multiplicities of infection (MOI), in the absence and presence of WASA-1 (Fig. 2a). This revealed that WonAB provides robust protection against ICP1 at low MOI when only a subset of cells is initially infected. In contrast, infection at high MOI, when most cells are simultaneously infected, resulted in a growth arrest (Fig. 2a). This phenotype is characteristic of abortive infection, wherein individual cell viability is sacrificed prior to the completion of phage replication to protect the surrounding bacterial population^46^. Indeed, time-lapse microscopy showed that while both WT and ΔWASA-1 cells swell upon infection, cells of the ΔWASA-1 go on to lyse within ∼20min, whereas WT cells remain in this swollen state without lysing for the 60min duration of the experiment (Fig. 2b, Extended Data Fig. 6 and movie 1). Moreover, these non-growing cells persisted in an intact state for several hours before eventually undergoing lysis (Extended Data Fig. 7a-b). During this time ICP1-infected cultures rebounded due to the selection of spontaneous O1-antigen mutants (Extended Data Fig. 7c), as has been previously described^7^. Importantly, this is likely an *in vitro* phenomenon since O1-antigen mutants are attenuated for virulence in animal infection models^47^.

**Fig. 2.**
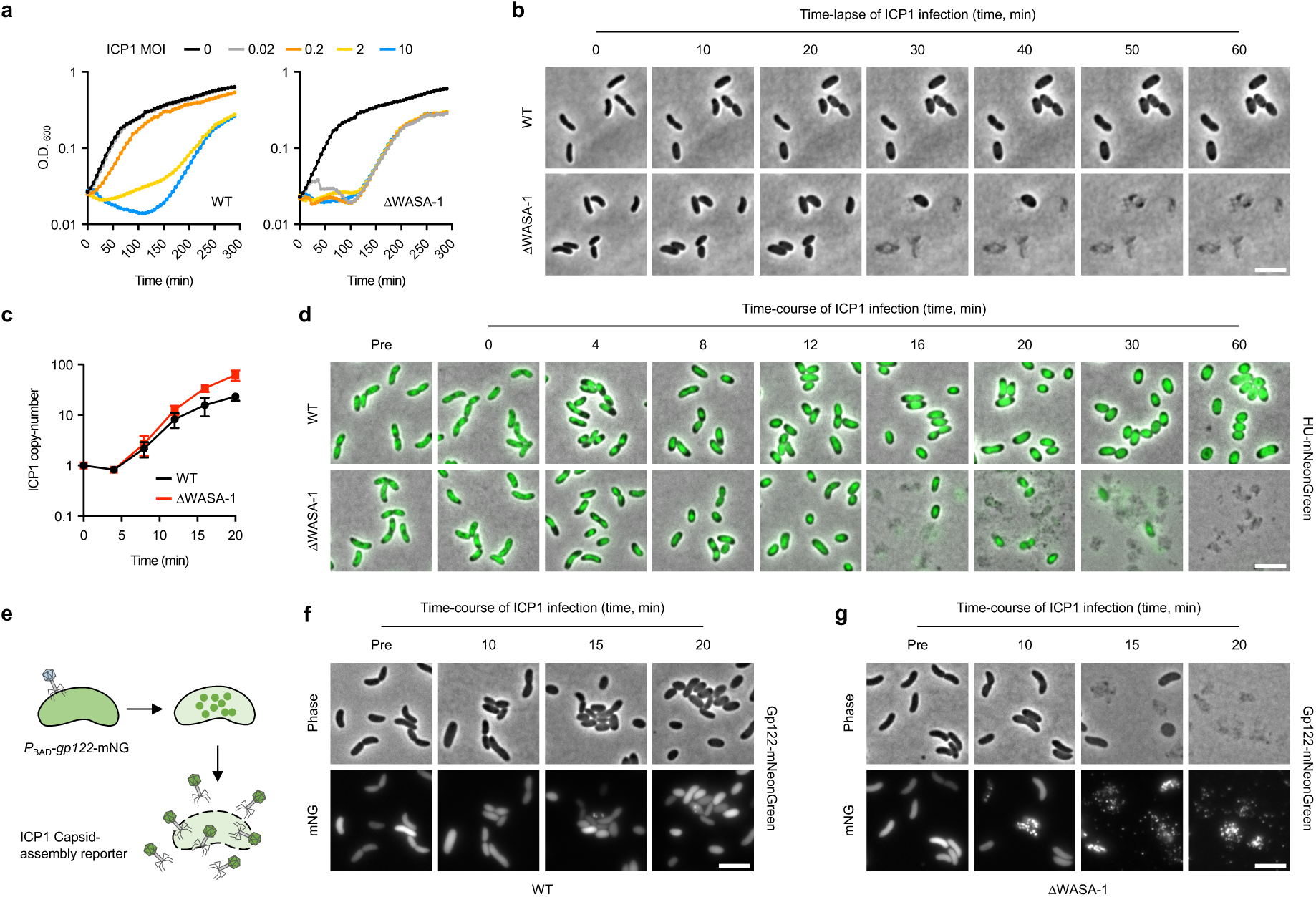
WonAB protects cells by lysis-independent abortive infection. (**a**) Growth kinetics of *V. cholerae* strain A1552 in the presence (WT) and absence of WASA-1 (ΔWASA-1), with either no phage or infected at time 0 with ICP1-2006 at the indicated MOI. Data are representative of three independent repeats. (**b**) Time-lapse microscopy comparing exponentially growing cells of *V. cholerae* A1552 WT and ΔWASA-1 strains after infection with ICP1-2006 at MOI 5. (**c**) Fold-change in ICP1-2006 genome copy number relative to time 0, as determined by qPCR, after infection of either WT or ΔWASA-1 strains with ICP1-2006 at MOI 0.1. (**d**) Time-course microscopy snapshots comparing cell morphology and cellular DNA content, as monitored by a HU-mNeonGreen fusion, in WT and ΔWASA-1 backgrounds following infection with ICP1-2006 at MOI 5. (**e-g**) Time-course microscopy snapshots comparing ICP1 capsid assembly, as monitored by the incorporation of a host-cell produced mNeonGreen (mNG) fusion to the ICP1 major capsid protein Gp122 (**e**), in either WT (**f**) or ΔWASA-1 (**g**) backgrounds, following infection with ICP1-2006 at MOI 5. All images are representative of the results of three independent experiments. Scale bars = 5 µm.

Efforts to isolate either a spontaneous or an evolved ICP1 escape mutant able to overcome WonAB have so far proved unsuccessful. Therefore, to study the mode of action of WonAB in more detail we investigated different stages of the ICP1 lifecycle in the absence and presence of WASA-1^8,48^. First, quantitative-PCR showed that ICP1 DNA replication proceeds similarly in both strain backgrounds, albeit with a 2.7-fold defect in the WT background, indicating that WonAB neither restricts ICP1 DNA on entry nor prevents the initiation of DNA replication (Fig. 2c). Second, microscopy revealed that upon infection the nucleoids of both strains undergo dramatic changes in morphology by 4min post-infection, with the nucleoid frequently appearing to contract along its long axis, likely representing the action of ICP1-encoded nucleases and host-cell takeover^48,49^ (Fig. 2d). Notably, whereas ΔWASA-1 cells then rapidly went on to lyse, DNA in WT cells persisted in a highly compacted state and went on to form toroidal structures, suggesting that WonAB does not act by simply degrading DNA. Third, microscopy of ICP1-infected cells producing a fluorescent protein fusion to the ICP1 major capsid protein revealed robust capsid assembly, lysis and release in the ΔWASA-1 background (Fig. 2e-g). In contrast, capsid assembly was largely absent in WT cells (Fig. 2f). Taken together, these results show that WonAB does not prevent the initial damage inflicted by ICP1, and suggest that WonAB is activated after the initiation of ICP1 DNA replication and prevents progression to structural protein assembly.

## WonAB activation shuts down cell growth by inhibiting translation

The data so far are consistent with WonAB aborting ICP1 infection via a lysis-independent mechanism that targets the host cell. To test whether WonAB can be artificially activated in the absence of infection, we overexpressed either the *wonAB* operon or each gene individually in an otherwise WT background (Fig. 3a and Extended Data Fig. 8a). While *wonAB* and *wonA* had no apparent phenotype, WonB overproduction was highly toxic, resulting in a growth arrest and a rapid loss in viability, with >99% of cells non-viable within 30min (Fig. 3a-c and Extended Data Fig. 8a). Furthermore, toxicity was dependent on WonB nuclease activity since the WonB[K71A] variant was non-toxic (Fig. 3b-c and Extended Data Fig. 8a). One explanation for these data could be that WonAB is a toxin-antitoxin system, with WonA acting as an antitoxin to neutralise the toxicity of WonB^50^. Indeed, the DarTG toxin-antitoxin system has recently been shown to have activity against ICP1^51^. Arguing against this hypothesis, however, WonB overproduction was not toxic in a Δ*wonAB* background, despite being produced at similar levels to the control (Fig. 3a and Extended Data Fig. 8b-c). Moreover, deletion of *wonA* or inactivation of WonA ATPase activity also abolished the toxicity associated with WonB overproduction (Fig. 3a and Extended Data Fig. 8b-c). Thus, these results rule out a simple sequestration mechanism and suggest that WonA is either required to activate WonB or that the two proteins act together.

**Fig. 3.**
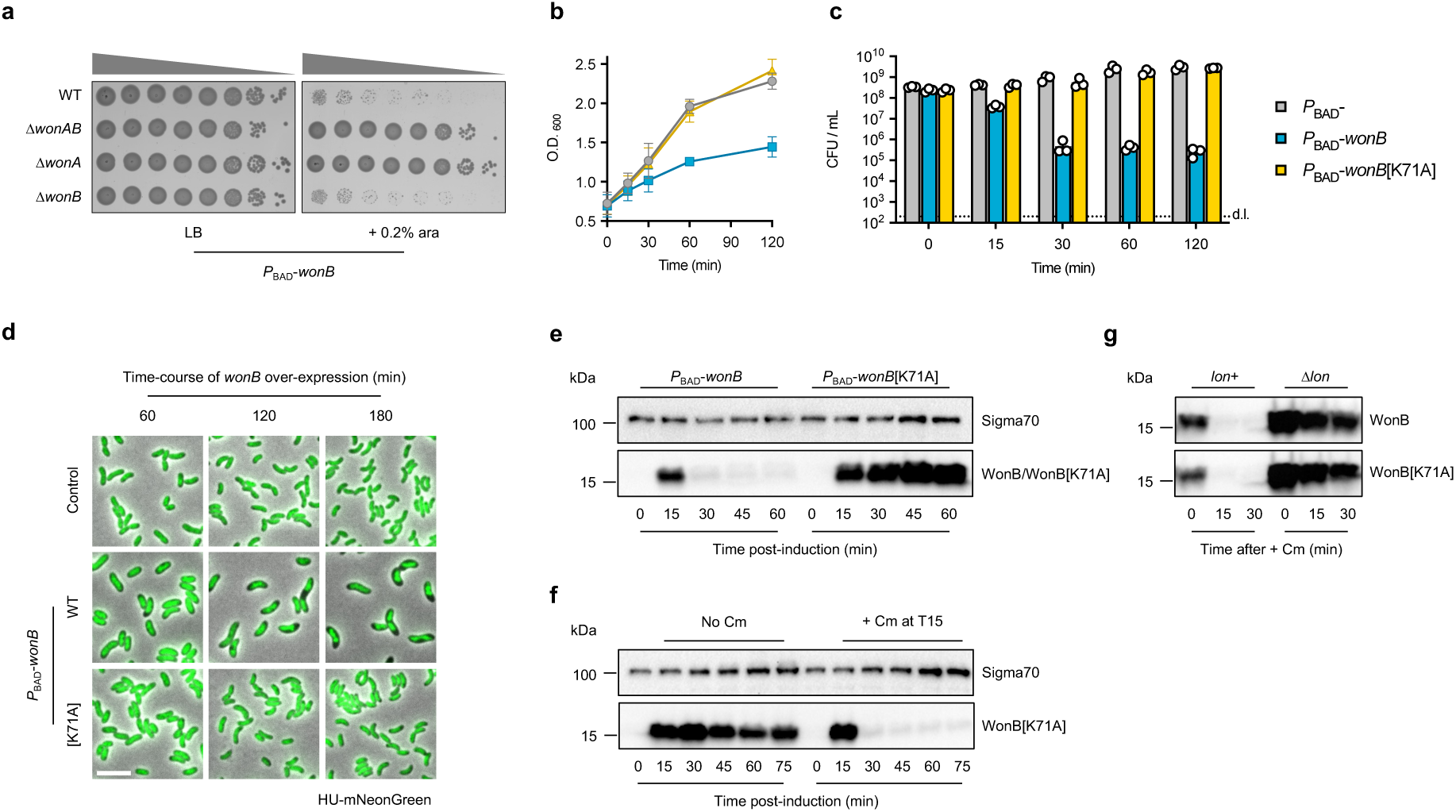
WonAB activation shuts down cell growth by inhibiting translation. (**a**) Toxicity assay evaluating the growth of 10-fold serial dilutions of *V. cholerae* strain A1552 cultures and the indicated derivatives on solid media, in the absence (LB) and presence (+ 0.2% ara) of WonB overproduction. (**b**-**c**) Toxicity assay evaluating the growth kinetics (**b**) and cell viability (**c**) of exponentially growing cultures in liquid media upon WonB overproduction following induction at time 0, as compared to a negative control strain and a strain overproducing the inactive WonB[K71A] variant. (**d**) Time-course microscopy snapshots showing the effect of WonB overproduction on cell morphology and cellular DNA content, as monitored by a HU-mNeonGreen fusion, compared to a negative control strain and a strain overproducing the inactive WonB[K71A] variant. Scale bar = 5 µm. (**e**) Western blot showing protein levels of WonB and WonB[K71A] over time following induction in exponentially growing cells at time 0. Predicted molecular mass of WonB, 16.1 kDa. Sigma70, was used as a loading control (**f**) Western blot showing protein levels of WonB[K71A] over time following induction in exponentially growing cells at time 0, in the absence and presence of chloramphenicol (Cm), added at 15min (T15). (**g**) Western blot showing protein levels of WonB and WonB[K71A] over time following chloramphenicol (Cm) addition, in the presence (*lon*+) and absence of Lon protease (Δ*lon*). Overexpression of genes in (**a**-**g**) was done from a chromosomally integrated transposon carrying the arabinose-inducible *P*_BAD_-promoter, induced by the addition of 0.2% arabinose. Charts show mean values, error bars show s.d. All images, charts and blots are representative of the results of three independent experiments.

Microscopy revealed that following WonB overproduction growth-arrested cells exhibit condensed nucleoids that at later stages display aberrant morphologies and are highly compacted (Fig. 3d and Extended Data Fig. 8d). Notably, the vast majority of cells retained DNA and purified genomic DNA also remained intact (Extended Data Fig. 8d-e), indicating that toxicity does not result from chromosomal DNA degradation. Since the nucleoid compaction phenotype resembled the effects of antibiotics that inhibit translation^52^, we investigated WonB production over time using Western blotting (Fig. 3e). Remarkably, this revealed that compared to the inactivated WonB[K71A] control, WonB is produced only for short time after induction, and by 30min post-induction was no longer detectable. Importantly, qRT-PCR showed that *wonB* mRNA levels were induced similarly between the two constructs and remained stable throughout the course of the experiment (Extended Data Fig. 8f). These data suggested that (i) WonB overproduction inhibits translation and (ii) that WonB is likely unstable. In agreement with these hypotheses, following inhibition of translation with chloramphenicol, WonB rapidly disappeared in a Lon protease-dependent manner (Fig. 3f-g), indicating that WonB is indeed unstable and subject to proteolytic degradation. Since in the absence of ICP1, WonB was toxic only when produced in excess of WonA, Lon-mediated degradation likely functions as a safety mechanism to reduce WonB accumulation and hence prevent auto-immunity. Collectively, these results support a model whereby upon activation WonAB aborts phage infection by inhibiting translation, and thus prevents the progression of the ICP1 lifecycle. Notably, although WonAB aborts the infection, cells are likely already irreversibly damaged by ICP1 during the initial stages of the infection. Furthermore, since the growth-arrest upon WonB overproduction does not result in genomic DNA degradation, these results suggest that the target of WonB nuclease activity could be a rRNA or tRNA.

## VSP-II^WASA^ carries two distinct anti-phage defence systems

The second genetic signature of the WASA lineage is a unique variant of VSP-II (Fig. 4a), wherein the chemotaxis gene cluster from the prototypical VSP-II is replaced by a set of three novel genes (A1552VC_00274-76) and a frame-shifted transposase gene^10,28,53^. Although these genes are not predicted to be part of a known defence system, and lacked activity against ICP1-3, bioinformatic analysis revealed that they encode proteins with domains characteristic of defence systems (see below). Consistent with this, when this cluster was expressed in *E. coli* we observed anti-phage activity against several major groups of the BASEL collection (Fig. 4b and Extended Data Fig. 9a). As outlined below, genetic dissection revealed that this cluster encodes two independent systems, with the two-gene operon VC_00274-75 encoding the <u>G</u>mrSD-like Type IV <u>R</u>Ease of <u>W</u>ASA strains, GrwAB (Fig. 4c-e) and VC_00276 encoding the *<u>V</u>. <u>c</u>holerae* <u>S</u>he<u>du</u>, *Vc*SduA (Fig. 4f-h).

**Fig. 4.**
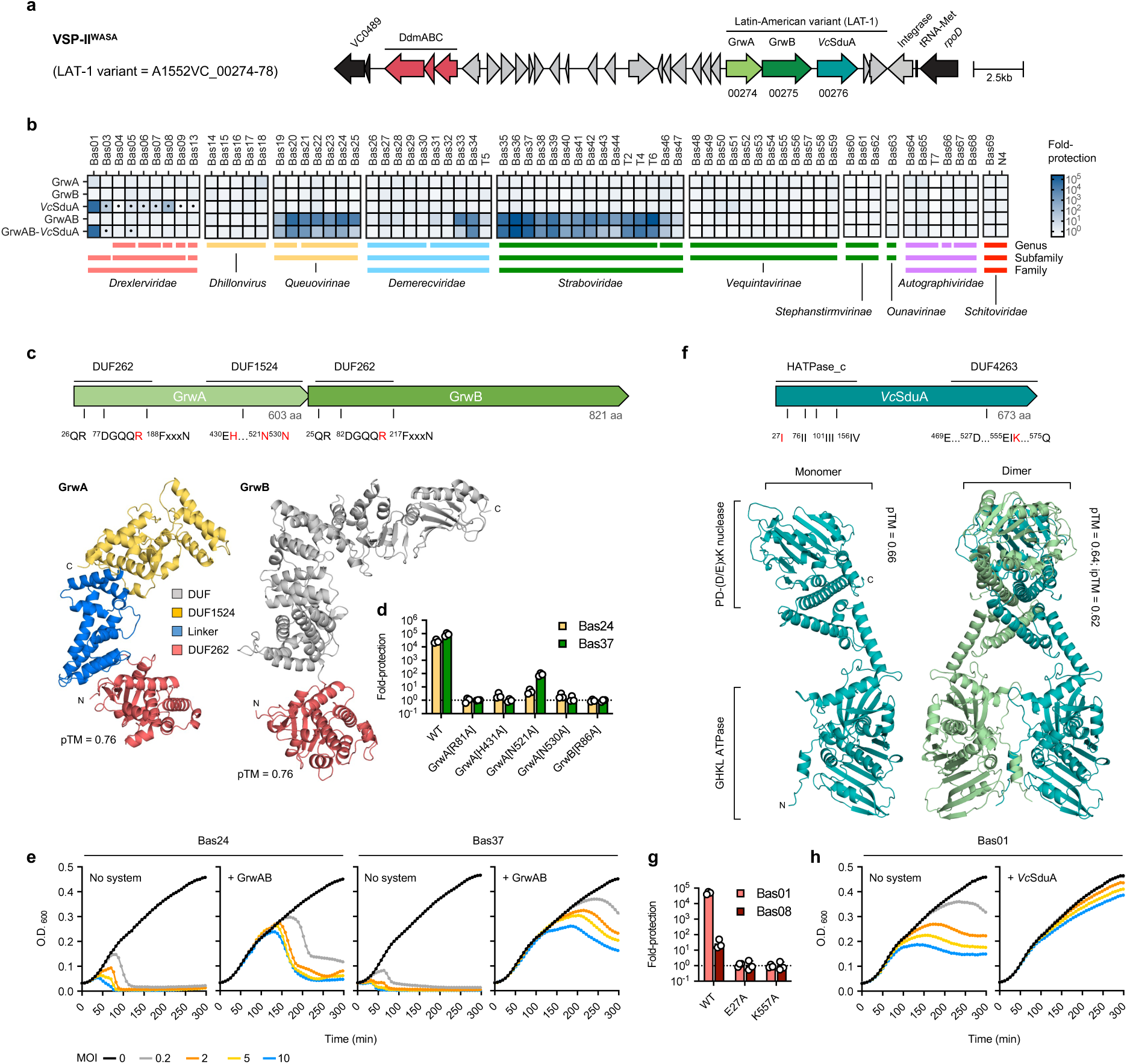
VSP-II^WASA^ variant encodes two distinct anti-phage defence systems. (**a**) Schematic of VSP-II^WASA^ highlighting the LAT-1 variant present in the WASA lineage and the previously characterised DdmABC system. (**b**) Fold-protection against *E. coli* phages of the BASEL collection conferred by the production of the indicated systems and components in *E. coli*, compared to a negative ‘no system’ control. Boxes with a filled dot indicate phages exhibiting altered plaque morphology. Data represent the mean of two independent repeats. (**c, f**) Predicted structures of GrwAB (**c**) and *Vc*SduA (**f**) generated using AlphaFold3 shown below schematics (top) indicating the conserved domains and motifs identified in each system. Residues targeted by site-directed mutagenesis are highlighted in red. (**d**, **g**) Fold-protection against the indicated *E. coli* phages conferred by site-directed variants of GrwAB (**d**) and *Vc*SduA (**g**) in *E. coli*, compared to a negative ‘no system’ control. Bar charts show the mean + s.d. of three independent repeats. (**e**, **h**) Growth kinetics of *E. coli* cultures in the absence (No system) and presence of either GrwAB (**e**) or *Vc*SduA (**h**), with either no phage or infected at time 0 with the indicated MOI. Data are representative of three independent repeats. Systems were expressed from a chromosomally integrated transposon using the arabinose-inducible *P*_BAD_-promoter. Growth media were supplemented with 0.2% arabinose.

## GrwAB is a two-component modification-dependent restriction system

GrwAB showed potent anti-phage activity against all members of the *Queuovirinae* and *Tevenvirinae* (*Straboviridae*), as well as some members of the *Markadamsvirinae* (*Demerecviridae*), and both genes were required for protection (Fig. 4b). Interestingly, the signature feature of the *Queuovirinae* and *Tevenvirinae* subfamilies is that their genomes contain hypermodified nucleobases with either 7-deazaguanine modified guanosines or sugar-modified hydroxymethyl cytosines, respectively, suggesting that GrwAB may recognise DNA modification^38,54,55^. In line with this hypothesis, GrwAB resembles the GmrSD family of modification-dependent restriction enzymes, such as GmrSD, BrxU and SspE^56–59^ (Fig. 4c and Extended Data Fig. 10a-d). These are typically single protein systems composed of an N-terminal DUF262 domain that likely functions as a DNA modification sensor, and that uses NTP binding/hydrolysis to regulate the activity of a C-terminal His-Me nuclease domain (DUF1524), which functions as an effector to target non-self DNA^56,58,60^. Indeed, GrwA is predicted to share this architecture, with an N-terminal DUF262 domain containing the highly conserved QR, DGQQR and FxxN motifs^56^, coupled by an α-helical linker to a C-terminal DUF1524 domain containing the canonical ββα fold and catalytic residues of the His-Me nuclease superfamily^61^ (Fig. 4c and Extended Data Fig. 10a-c). In contrast, and unlike known GmrSD systems, GrwB encoded by the second gene contains an N-terminal DUF262 domain followed by multiple domains of unknown function (Fig. 4c and Extended Data Fig. 10d). Notably, variants designed to disrupt either the NTP hydrolysis of DUF262 in GrwA[R81A] and GrwB[R86A], or the nuclease activity of DUF1524 in GrwA[H431A]/[N521A/[N530A] all abolished anti-phage activity (Fig. 4d), indicating that GrwA and GrwB function together. Indeed, structural modelling of a potential GrwAB complex supports the idea that the two proteins interact (Extended Data Fig. 10e).

Importantly, multiple independent escaper mutants of Bas24 were obtained that overcome defence by GrwAB (Extended Data Fig. 11a). These escapers all contain changes in either the active site or the DNA-binding surface of DpdA – the transglycosylase that inserts the 7-deazaguanine modification into the DNA – consistent with the disruption of DNA modification leading to immune evasion^62^ (Extended Data Fig. 11b-d). Interestingly, growth kinetics of phage-infected cultures producing GrwAB lacked the growth arrest or premature lysis typical of abortive infection (Fig. 4e). However, cultures did go on to exhibit moderate-to-severe lysis, which microscopy revealed was due to the presence of non-dividing filamentous cells (Extended Data Fig. 12 and movie 2), suggesting that either GrwAB is unable to clear the infection prior to the cell sustaining irreversible damage or that the cell is subject to collateral damage during phage defence. Finally, bioinformatic analyses revealed that homologues of GrwAB are found in ∼1% of genomes examined (Extended Data Fig. 10f and Supplementary Table 4), suggesting that GrwAB represents a new member of an emerging family of two-gene GmrSD-like modification-dependent restriction enzymes that includes the recently discovered TgvAB system present in *V. cholerae* VPI-2^25,26^. Similar to TgvAB, the requirement for both GrwA and GrwB suggests that these proteins likely act as a complex, which could represent either a novel mode of action or else a mechanism to overcome phage-encoded anti-defences. Notably, despite the similarities, GrwAB has an expanded range compared to TgvAB and is also able to recognise 7-deazaguanine modified DNA. Furthermore, although GrwA, TgvB and GmrSD all share a similar domain architecture, except for the shared N-terminal DUF262 domain GrwB and TgvA are distinct from one another^25^.

## *Vc*SduA is a Shedu system with a GHKL ATPase N-terminal domain

*Vc*SduA showed activity against all members of the T1-like *Drexlerviridae* family, with strong protection against Bas1 (>10,000-fold), Bas03 and Bas08 (10-100-fold), while the remainder showed a reduced plaque diameter that in the case of Bas03 and Bas05 was drastic (Fig. 4b and Extended Data Fig. 9b). Sequence analysis and structural modelling showed that *Vc*SduA encodes an N-terminal GHKL (Gyrase, Hsp90, Histidine Kinase, MutL) ATPase domain (HATPase_c) containing the highly conserved motifs I-IV that define this ATPase superfamily^63,64^, followed by a C-terminal PD-(D/E)xK nuclease domain (DUF4263) that is also found in the Shedu defence system^20,65,66^ (Fig. 4f and Extended Data Fig. 13a-c). Variants designed to disrupt either ATP hydrolysis *Vc*SduA[E27A] or nuclease activity *Vc*SduA[K557A] both abolished anti-phage activity despite being produced at similar levels to WT control (Fig. 4g and Extended Data Fig. 13d). Furthermore, growth kinetic experiments with phage-infected cultures revealed similarly robust protection at all tested MOIs (Fig. 4h), consistent with *Vc*SduA targeting and restricting the phage directly while preserving the viability of the host cell.

Proteins with a similar domain organisation to *Vc*SduA were previously annotated as paraMORC3, and have recently been proposed to be part of the Shedu defence system family due to the presence of the signature Shedu domain DUF4263^64,65^. Indeed, Shedu has recently been re-classified as consisting of a shared nuclease domain with diverse N-terminal regulatory domains (Type I), including various enzymatic domains (Type I-D)^65^. Strikingly, modelling of *Vc*SduA as a dimer (Fig. 4f), which are typical for GHKL ATPases, confidently predicted an entwined configuration with the GHKL ATPase domain sitting below a cavity formed by a clamp-like middle domain, which in other GHKL family members mediates binding to DNA and protein substrates^63,67^. We therefore hypothesise that the GHKL ATPase domain functions to control the activity of the C-terminal nuclease domain. Interestingly, modelling of *Vc*SduA as a tetramer was not well-supported (Extended Data Fig. 13e). Furthermore, *Vc*SduA homologues were present in ∼1.2% of genomes examined (Extended Data Fig. 13f and Supplementary Table 5), though GrwAB and *Vc*SduA were rarely found together outside of *V. cholerae*, consistent with the demonstration above that they are independent systems. Overall, these data show that *Vc*SduA is a newly identified member of the Shedu family and the first validated member of the type I-D class.

## *Vc*SduA protects *V. cholerae* against vibriophage X29

Next, to determine whether GrwAB and *Vc*SduA were functional in their native *V. cholerae* host, we first searched the NCBI virus database for vibriophages with modified genomes that might be targeted by GrwAB (see methods). However, no such phage targeting *V. cholerae* were identified (Supplementary Table 6). We therefore screened a collection of phages from the Félix d’Hérelle Reference Center for Bacterial Viruses, focussing on phage X29^68^ as we determined that it requires the O-antigen for infection (Fig. 5a). Importantly, Darracq *et al.*, recently showed that the VSP-II encoded DdmABC defence system^24^ protects *V. cholerae* against X29^69^. Indeed, deletion of *ddmABC* in the non-WASA 7PET strains tested here was sufficient to fully restore X29 plaque formation (Fig. 5b). In contrast, deletion of *ddmABC* in the WASA-lineage strains A1552 and C6706 had no effect (Fig. 5b), raising the possibility that the LAT-1 encoded defence systems of VSP-II^WASA^ were active against X29. Consistent with this idea, while deletion of *grwAB* or *VcSduA* alone had no effect, deletion of either the entire LAT-1 region or *VcSduA* alone in a Δ*ddmABC* background restored X29 plaque formation to levels comparable to that of a complete VSP-II deletion (Fig. 5c). Furthermore, ectopic expression of *VcSduA* complemented the Δ*ddmABC*Δ*VcSduA* deletion and conferred complete protection against X29 in non-WASA strains deleted for *ddmABC* (Fig. 5d-e). Collectively, these data show that *Vc*SduA is active in its native host organism at native expression levels, and show that WASA lineage strains have two distinct mechanisms to defend against phage X29.

**Fig. 5.**
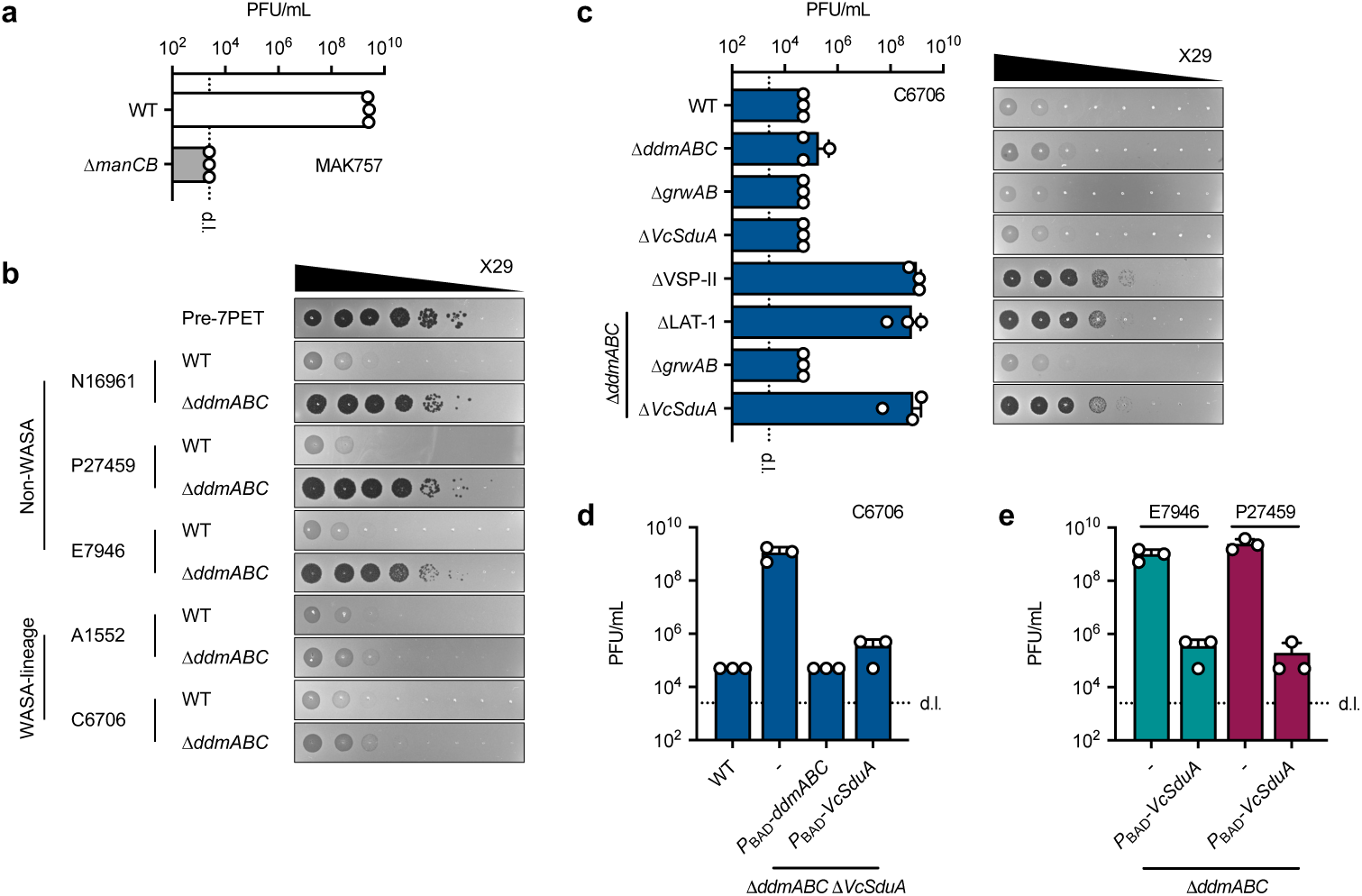
*Vc*SduA protects *V. cholerae* against *vibrio* phage X29. (**a**) Plaque assay showing X29 plaque-forming units (PFU/mL) on the pre-7^th^ pandemic strain MAK757, in the presence and absence (Δ*manCB*) of the O1-antigen. (**b**) 10-fold serial dilution plaque assays of X29 on wave 1 *V. cholerae* strains, comparing the effect of deleting *ddmABC* in non-WASA (N16961, P27459, E7946) and WASA-lineage (A1552, C6706) strains, below a pre-7^th^ pandemic control (MAK757). (**c**) Plaque assay showing X29 PFU/mL obtained on C6706 derivates deleted for combinations of *ddmABC* with ΔLAT1 (Δ*grwAB*-*VcSduA*), Δ*grwAB* and Δ*VcSduA*, alongside representative images of each 10-fold serial dilution plaque assay. Note that strain C6706 was used in these assays because the rifampicin antibiotic resistance marker in strain A1552 partially interferes with X29 infection. (**d-e**) Plaque assays testing the ability of *Vc*SduA production to complement X29 protection in a C6706Δ*ddmABC*Δ*VcSduA* background (**d**) and in Δ*ddmABC* backgrounds of the non-WASA linage strains E7946 and P27459 (**e**). Bar charts show the mean + s.d. from 3 independent repeats. d.l. = detection limit. Images are representative of the results of three independent experiments.

## Discussion

The main discovery of this work is that the two unique genetic signatures (WASA-1 and VSP-II^WASA^) of the WASA lineage encode three novel antiviral defence systems: WonAB, GrwAB, *Vc*SduA. Previously, these elements have primarily been used as a molecular signature to track the origins of these strains^10,28–30,70^, but their significance for the success of the lineage and the Latin American cholera epidemic has remained unknown. Here we demonstrate that combined these systems have broad activity against diverse bacteriophage families and target two-thirds of the isolates in the BASEL bacteriophage collection, including those with highly modified genomes. Importantly, we also show that in the native *V. cholerae* host, the WonAB system carried on the WASA-1 prophage renders strains resistant to ICP1 - the major predatory phage of pandemic *V. cholerae*^8^. Given the ongoing arms race between *V. cholerae* and ICP1^11,14^ and the suggested role of ICP1 predation in cholera epidemics^3,4,6,7^, it is therefore reasonable to assume that this would have provided a significant selective advantage to the lineage. Nevertheless, since WonAB can also target other phages, then together with the systems carried on VSP-II^WASA^, this selective advantage may have been against phages other than ICP1. Indeed, the VSP-II^WASA^-encoded *Vc*SduA system protects WASA lineage strains against vibriophage X29, a myovirus that was originally isolated from a cholera patient in India in 1927^68,71^. Notably, spacers targeting X29 are abundant in the CRISPR-Cas systems present in classical 6^th^ pandemic and non-7PET strains^72–74^, suggesting that X29 (or related phages) may be commonly encountered. Interestingly, since *Vc*SduA shares redundancy with DdmABC it could potentially serve to protect against X29 variants that have evolved DdmABC-resistance. However, given *Vc*SduA activity against *Drexlerviridae* in *E. coli* it is equally possible that it functions primarily against other non-DdmABC sensitive phages.

Prior to the start of the epidemic in 1991, Latin America had been free from pandemic cholera for almost 100 years^27,75^. However, following the introduction of the WASA lineage from West Africa in the late 1980’s, an explosive cholera epidemic spread rapidly throughout Latin America, with >500,000 cases in Peru alone in 1991-92, and ongoing outbreaks continuing until 2001^10,27,30,31,75,76^. Given our findings we therefore propose that the combined activities of the defence systems encoded on WASA-1 and VSP-II^WASA^ directly contributed to the success of the WASA lineage and envisage at least two possible scenarios by which this could have occurred. Firstly, the acquisition of these systems in Africa could have allowed the strains to resist local vibriophages, triggering the proliferation of the lineage that therefore facilitated the transfer to South America. Notably, cholera outbreaks were ongoing in Western and Central Africa just prior to the start of the Latin American epidemic, and intensified in the years afterwards^30,31^. Furthermore, strains containing WASA-1 and VSP-II^WASA^ continued to circulate in Western/Central Africa up until the late 1990’s, including in the strains responsible for the 1987-1996 Angolan cholera epidemic^30,31,77^. Remarkably, we found WASA-1 (99.7% identity to A1552) in a recently sequenced non-7PET *V. cholerae* strain isolated in Switzerland in 2019 from a patient with a travel history to Morocco^78^, suggesting that WonAB-containing WASA-1 continues to circulate in Africa. Secondly, the defences provided by WASA-1 and VSP-II^WASA^ could have played a more direct role in the transmission, for example by allowing the WASA lineage to overcome an entry barrier posed by phages associated with local non-pandemic *V. cholerae* strains endemic to South America that might have prevented previous introductions. In support of these proposals, the ability to defend against phage predation is thought to play a key role in the observed evolution of local *V. cholerae* lineages, which then go on to be successful before subsequently being replaced^11,14^. Indeed, the continuous antagonistic co-evolution of bacterial defences and phage encoded anti-defences drives the gain and loss of defence systems in natural *vibrio* populations, with such defences often encoded on mobile genetic elements (MGE) and accounting for a large proportion of the accessory genome^18,79^. Thus, while 7PET strains are typically considered to be clonal based on the phylogeny of the core genome, closely related-strains can harbour diverse MGEs encoding defence systems that vary over time in response to the dominant phages^11,14^. This, together with the results presented here, reinforces the need to consider the role of the accessory genomes in the success of novel lineages and therefore understanding their pandemic potential. Finally, while further work will be necessary to decipher the mechanistic details of the newly discovered defence systems, this work provides new insights into the evolution of one of the major epidemic lineages of the seventh cholera pandemic.

## Materials and Methods

### Bacterial strains, plasmids and bacteriophages

The bacterial strains, plasmids, bacteriophages and defence systems used in this work are detailed in Supplementary Table 7. The primary *V. cholerae* strain used throughout this work, A1552, is an O1 El Tor (Inaba) strain isolated from a patient in 1992 following an outbreak of cholera linked to a commercial airline flight between Lima, Peru and Los Angeles, California, U.S.A^81–83^. The primary *E. coli* strain used throughout this work is the non-arabinose metabolising strain MG1655Δ*araCBAD*^84^. *E. coli* strains S17-1λ*pir* and MFD*pir* were used for cloning and bacterial mating with plasmids containing the conditional R6K origin of replication^85,86^.

### Growth conditions

Bacteria were cultured either in Lysogeny broth (LB-Miller; 10 g/L tryptone, 5 g/L yeast extract, 10 g/L NaCl; Carl Roth, Switzerland) with shaking at 180rpm or on LB + 1.5% agar plates, at 37°C. Transformation of *V. cholerae* was done on chitin powder (Alfa Aesar via Thermo Fisher, U.S.A) suspended in half-concentrated Instant Ocean medium (Aquarium Systems) and sterilised by autoclaving. Thiosulfate citrate bile salts sucrose (TCBS) agar plates were used for counter-selection against *E. coli* following bacterial mating (Sigma-Aldrich). SacB-based counter-selection for allelic exchange was done on NaCl-free media containing 10% sucrose. Counter-selection of *pheS*\*-carrying strains (see below) was done on LB agar plates supplemented with 20 mM 4-chloro-phenylalanine (cPhe). Ampicillin (Amp; 100 μg/mL), chloramphenicol (Cm; 2.5 µg/mL), gentamicin (Gent; 25-50 μg/mL), kanamycin (Kan; 75 μg/mL), streptomycin (Strep; 100 μg/mL) and rifampicin (Rif; 100 μg/mL) were used for selection, as required. Growth of MFD*pir* was supported by the addition of 0.3 mM diaminopimelic acid (DAP; Sigma-Aldrich). Unless stated otherwise, genes under control of the *P*_BAD_-promoter were induced by supplementing growth media with 0.2% L-arabinose.

### Strain construction

Genetic engineering of *V. cholerae* was done using natural competence for transformation to introduce deletions marked with antibiotic resistance genes^87^, which when necessary were removed using FLP recombination (TransFLP;^88,89^). Alternatively, antibiotic cassettes linked to the modified phenylalanyl-tRNA synthetase *pheS*\* (*pheS*[A294G/T251A]) were removed by natural transformation using the Trans2 method^90^. Scar-less, marker-less modifications were introduced using allelic exchange with derivatives of the counter-selectable plasmid pGP704-Sac28^91^. A mini-Tn7 transposon carrying *araC* and the indicated gene(s) of interest under control of the *P*_BAD_-promoter was integrated downstream of the *glmS* locus in *V. cholerae* and *E. coli* by introducing derivatives of pGP704-TnAraC via tri-parental mating, as previously described^92^. Plasmids were constructed using either standard restriction enzyme cloning or by inverse PCR, and were used exclusively as intermediates for strain construction. All strains and constructs were verified by PCR, and either Sanger sequencing or Oxford Nanopore Technologies (ONT)-based full plasmid sequencing (Microsynth AG, Switzerland).

### *V. cholerae* bacteriophage methods

Vibriophages were propagated on *V. cholerae* strains E7946 or MAK757 (ICP1 and ICP3) and E7946Δ*manCB* (ICP2). Briefly, overnight cultures were back-diluted and grown in LB at 37°C, 180rpm to exponential phase (optical density at 600nm, OD_600_ *c.a.* 0.5-0.6), infected with the relevant phage at a multiplicity of infection (MOI) of <0.01. Following lysis of the culture, debris was removed by centrifugation (10min; 4,000 x *g*; RT), lysates passed through a 0.2µm filter, and stored at 4°C after the addition of chloroform to 1%. Phage titres were determined by plaque assays on strain E7946 using the small-drop plaque assay on double-layer plates, as described below. To quantify ICP and X29 phage infections, 100µL and 10µL, respectively, of overnight culture was added to molten top-agar (LB + 0.5% agar), poured on top of a bottom agar (LB + 1.5% agar), and allowed to solidify at RT. For complementation of *wonAB* and *VcSduA*, cultures were grown either for 2h + 0.2% arabinose or overnight with 0.02% arabinose, respectively, and the top agar was also supplemented with arabinose at the respective concentration. Phages were serially-diluted (10^−1^-10^−8^), 4µL of each dilution spotted onto plates, and allowed to dry under a laminar flow hood. Plates were imaged after overnight incubation at 37°C, and the plaque-forming units (PFU/mL) enumerated. The theoretical detection limit in this experiment was 2.5×10^3^ PFU/mL. In cases where no discrete plaques were observed but a zone of non-specific lysis was apparent (*e.g.* ICP1 on strain A1552; Fig. 1a), the last dilution exhibiting this phenotype was counted as 20 plaques. To determine whether these zones of non-specific lysis resulted from phage propagation or “lysis from without” (*i.e.* lysis at high phage concentration without viable phage production) re-streak tests were performed, as previously described^18^. Material from the highest phage concentration spot on plaque assay plates was re-streaked onto freshly poured top agar plates containing the ICP1-sensitive strain A1552ΔWASA-1.

### *E. coli* bacteriophage methods

The BASEL (BActeriophage SElection for your Laboratory) collection of *E. coli* bacteriophages was obtained from A. Harms, ETH Zürich^38^. Infections were quantified using the small-drop plaque assay described above, with the following modifications. Briefly, overnight cultures of *E. coli* MG1655Δ*araCBAD* were back-diluted 1:100 in 3mL LB + 5mM CaCl_2_, 20mM MgSO_4_, 0.2% arabinose and grown 2h at 37°C, 180rpm to reach exponential phase, before being diluted 1:40 in molten top agar, supplemented with 5mM CaCl_2_, 20mM MgSO_4_ and 0.2% arabinose. Phages were serially-diluted (10^−1^-10^−8^), 5µL of each dilution spotted onto plates, and allowed to dry under a laminar flow hood. Plates were imaged after overnight incubation at 37°C, and the plaque-forming units (PFU/mL) enumerated. The theoretical detection limit in this experiment was 2×10^3^ PFU/mL. Fold-protection was calculated as the ratio of PFU/mL on the strain of interest to that of the no system control strain *E. coli* MG1655Δ*araCBAD*-TnAraC.

### Quantification of infectious ICP1 phages

To monitor the production of infectious ICP1 phages, overnight cultures of the *V. cholerae* strains indicated in the text were back-diluted 1:100 in 2.5mL LB in 14mL test-tubes and grown at 37°C, 180rpm to exponential phase (OD_600_ *c.a.* 0.5-0.6) before being infected with *c.a.* 10^3^-10^4^ PFU/mL of ICP1-2006. Infected cultures were incubated at 37°C, 180rpm and sampled at 30min intervals for 2.5h. At each time-point, 300µL culture was mixed with 30µL of chloroform, vortexed vigorously for 10s and centrifuged (3min; 20,000 x *g*; RT). To determine the starting titre at time 0 samples were prepared from LB only controls. Chloroform-free supernatants were transferred to new tubes, serially diluted (10^0^-10^−7^) and PFU/mL enumerated by a plaque assay on plates seeded with the ICP1-sensitive *V. cholerae* strain E7946. The theoretical detection limit in this experiment was 2.5×10^2^ PFU/mL.

### Bacteriophage infection kinetics

To monitor the kinetics of ICP1 infection, overnight cultures of the *V. cholerae* strains indicated in the text were back-diluted 1:100 in 2.5mL LB in 14mL test-tubes and grown at 37°C, 180rpm for 1h45min to exponential phase (OD_600_ *c.a.* 0.5-0.6). 180µL aliquots of cultures diluted 1:5 in pre-warmed LB were added to the wells of a 96-well plate containing 20µL of ICP1-2006 at the indicated MOI (in technical triplicate). Bacterial growth (OD_600_) at 37°C with shaking was followed using a SpectraMax i3x (Molecular Devices) plate reader at 6-min intervals for a total of 49 cycles. The kinetics of Bas01, Bas24 and Bas37 infection in *E. coli* were monitored exactly as described above, except that *E. coli* strains were grown for 2h in LB + 5mM CaCl_2_, 20mM MgSO_4_, 0.2% arabinose and aliquots were diluted 1:10. *E. coli* phages used for plate reader experiments were propagated on MG1655Δ*araCBAD*, as described above.

### Microscopy

Imaging was conducted using Zeiss Axio Imager M2 and Zeiss Axio Observer Z1 epifluorescence microscopes, equipped with AxioCam MRm cameras and controlled by Zeiss Zen software (v.2.6 blue edition). Images were captured using a Plan-Apochromat 100x/1.4-NA Ph3 oil objective, and for fluorescence microscopy illumination with an HXP120 lamp. For snap-shot imaging, cells were immobilized on slides coated with 1.2% (wt/vol) agarose in PBS, covered with a No. 1 coverslip. For time-lapse microscopy, cells were immobilized on slides coated with 1.2% (wt/vol) agarose in either LB (ICP1 infection of *V. cholerae*) or LB + 5mM CaCl_2_, 20mM MgSO_4_, 0.2% arabinose (Bas24 infection of *E. coli*) and imaged automatically at the indicated time-points, within a temperature-controlled stage-top chamber (H301-K-Frame; OKO Lab, Italy) set to 37°C with an accompanying objective-heater. Images were prepared for publication using ImageJ (version 2.1.0/1.53h; imagej.net/software/fiji), and where needed, drift was corrected using the HyperStackRegPlus function of MicrobeJ^93^.

### Time-lapse and time-course microscopy of ICP1 infection

To image ICP1 infection in *V. cholerae*, overnight cultures of the strains indicated in the text were back-diluted 1:100 in 2.5mL LB in 14mL test-tubes and grown at 37°C, 180rpm for 1h45min to exponential phase (OD_600_ *c.a.* 0.5-0.6). For time-lapse microscopy, an aliquot of the culture was then mixed with ICP1-2006 to achieve a MOI of 5, and immediately transferred to a slide and imaged, as described above. For time-course microscopy, 180µL aliquots of cultures were mixed with 20µL ICP1-2006 in 2mL microcentrifuge tubes to achieve a MOI of 5 and incubated flat at 37°C, 180rpm. At the indicated time-points, tubes were removed from the incubator and samples immediately transferred to a slide and imaged, as described above. To monitor cellular DNA content, strains were used encoding a C-terminal mNeonGreen fusion to the Histone-like DNA-binding protein HU (HU-mNeonGreen) separated by a GSGSGS linker, and expressed from the native *hupA* locus. To monitor ICP1 capsid assembly, infections were performed in strains with a chromosomally integrated transposon carrying a C-terminal mNeonGreen fusion to the ICP1 major capsid protein Gp122 (Gp122-mNeonGreen), separated by a GSGSGS linker, and expressed under the control of the arabinose-inducible *P*_BAD_-promoter. The fusion was induced by the inclusion of 0.02% arabinose in the growth media.

### Time-lapse microscopy of Bas24 infection

To image Bas24 infection in *E. coli*, overnight cultures were back-diluted 1:100 in 3mL LB + 5mM CaCl_2_, 20mM MgSO_4_, 0.2% arabinose in 14mL test-tubes and grown at 37°C, 180rpm for 2h to mirror the conditions used for the BASEL screen above. An aliquot of the culture was then mixed with Bas24 to achieve a MOI of 10, and allowed to stand at RT for 2min before being transferred to a slide and imaged.

### Quantitative PCR (qPCR) of ICP1 genome replication

ICP1 genome replication was quantified by quantitative PCR (qPCR) using a protocol modified from^13^. Briefly, overnight cultures were back-diluted 1:100 in 2.5mL LB in 14mL test-tubes and grown at 37°C, 180rpm for 1h45min to exponential phase (OD_600_ *c.a.* 0.5-0.6). ICP1-2006 was then added to a MOI of 0.1, tubes briefly vortexed to ensure proper mixing, and a 20µL sample immediately withdrawn (*t* = 0) and heat-inactivated at 95°C for 20min. Sampling was repeated following incubation at 37°C, 180rpm for 4, 8, 12, 16 and 20min post-infection. Samples were analysed using a LightCycler 96 Instrument (Roche) with SYBR Green I (Fast Start Essential DNA Green Master; Roche) to detect amplification products using the ICP1-specific primers qICP12006E_148-f (ACTTTGGTGCGTGAAGAAGG) and qICP12006E_148-r (ACTTGCTCACCTGAATGGTC). ICP1 genome copy number levels are presented relative to *t* = 0, and were analysed with LightCycler 96 software v.1.1.0.1320 (Roche) with the absolute quantification method using a standard curve of purified ICP1-2006 genomic DNA.

### WonB toxicity assay

Overnight cultures were back-diluted to OD_600_ 0.05 in LB and grown 1h45min at 37°C, 180rpm to exponential phase (OD_600_ *c.a.* 0.5-0.6), induced by the addition of 0.2% arabinose (*t* = 0), and incubation continued for 120min. The OD_600_ of each culture was measured at *t* = 0, 15, 30, 60, and 120min. At each time point, samples were collected and processed to determine the (i) CFU/mL by serial-dilution and plating on LB agar, (ii) transcript levels via qRT-PCR and (iii) genomic DNA was prepared using a GenElute Bacterial Genomic DNA Kit (NA2110; Sigma-Aldrich) according to the manufacturer’s instructions. Samples to determine protein levels by Western blotting were collected at the time-points indicated in the text and processed as described below. To determine the stability of WonB protein levels over time, translation was inhibited by the addition of chloramphenicol to 200 µg/mL.

### Quantitative reverse transcription PCR (qRT-PCR) of *wonAB* transcript levels

2mL culture samples from the WonB toxicity assay (above) were harvested by centrifugation (3min; 20,000 x *g*; 4°C), cell pellets resuspended in Tri-Reagent (Sigma-Aldrich), snap-frozen in a dry ice-ethanol bath, and stored at −80°C. RNA extraction, cDNA synthesis, and qPCR was then performed exactly as previously described^94^, using a LightCycler 96 Instrument (Roche). Transcript levels are presented relative to mRNA levels of the reference gene *gyrA*, and were analysed with LightCycler 96 software v.1.1.0.1320 (Roche) using the standard curve method. Primers used to detect *wonA and wonB* were: qVC_01233-F (GTCCACAAGCAAACGGTAAG) and qVC_01233-R (TTCTCCCATGAATACCTCGG); qVC_01234-F (ACAACTGTCTCAAAGGATCC) and qVC_01234-R (GCTCAATATCCAGTCACATC).

### Western Blotting

Unless stated otherwise, overnight cultures were back-diluted 1:100 and grown at 37°C, 180rpm for 3h. Lysates were prepared by resuspending pelleted bacteria in 2x Laemmli buffer (Sigma-Aldrich), normalised to optical density (100 μL buffer per OD unit), and samples were then heated at 95°C, 15min. Proteins were resolved on Mini-PROTEAN TGX Stain-Free precast gels (10% or 12%, as required; Bio-Rad) and transferred onto PVDF membranes using a Trans-Blot Turbo Transfer System (Bio-Rad) according to the manufacturer’s instructions. Membranes were blocked in 2.5% skim milk in TBST (1x Tris-buffered saline with 0.1% Tween-20) with agitation at either RT for 1h or at 4°C overnight. Primary antibodies were added at a dilution of 1:500 in TBST and membranes incubated at RT for 1h. Membranes were then washed three times with TBST before being incubated at RT for 1h with anti-rabbit IgG conjugated to HRP (A9169, Sigma-Aldrich) diluted 1:20,000 in TBST. Membranes were washed as above and visualised by chemiluminescence using Lumi-Light^PLUS^ Western Blotting Substrate (Roche). Primary antibodies against WonA (2210455), WonB (2210453) and *Vc*SduA (2310059) were custom-raised in rabbits against synthetic peptides (Eurogentec), and their specificity validated by comparing strains lacking the relevant proteins. Uniform sample loading was verified by the intensity of non-specific bands on the membranes or by detection of Sigma70 (1:10,000 diluted anti-Sigma70-HRP antibody from BioLegend, U.S.A).

### Identification of defence systems, protein domains, motifs and structural predictions

The repertoire of known defence systems present in *V. cholerae* strains was identified using DefenseFinder (v.1.2.4) and PADLOC (v.2.0.0) (accessed May 14^th^ 2024), see Supplementary Table 1 for details^95,96^. Conserved protein domains were initially identified by searching the PFAM and NCBI Conserved Domain Database with the MOTIF Search tool (genome.jp/tools/motif/; accessed July 22^nd^ 2024), and was supplemented by remote homology searches using HHpred (PDB_mmCIF70_8_Mar, default database; accessed July 22^nd^ 2024), and structural alignments performed with DALI using structural predictions generated by AlphaFold3^97–99^. The predicted template modelling (pTM) score, and where appropriate, the interface predicted template modelling (ipTM) score, are shown alongside each model^99^. For full confidence metrics for AlphaFold models, see Supplementary Fig. 1. Sequence logos showing the conservation of motifs in each system were generated using Weblogo 3 from multiple sequence alignments done using MAFFT^100,101^.

### Identification and distribution of WASA-1

A BLASTN search of the *V. cholerae* A1552 WASA-1 sequence (default parameters; evalue 1e-10) against 45,725 bacterial genomes (NCBI, Refseq database, complete and chromosome level genomes, downloaded on May 22^nd^ 2024) revealed hits exclusively in the *Vibrio* genus. The BLASTN search was therefore repeated against a database consisting of 7,624 genomes, covering all available assemblies within the *Vibrio* genus (Refseq database, genomes from contig to complete level assemblies, downloaded on May 30^th^ 2024). A second complementary BLASTN search was also performed with the additional max_hsps (high-scoring pair) parameter set to 1. Initial hits were then manually curated to identify complete WASA-1 assemblies, as shown in Supplementary Table 2. WASA-1 conservation was visualised using clinker (v.0.0.28, default parameters)^102^.

### Identification and distribution of defence systems

The presence of the WonAB, OLD-ABC ATPase + Novel REase, GrwAB and *Vc*SduA defence systems was determined against 45,725 bacterial genomes (NCBI, Refseq database, complete and chromosome level genomes, downloaded on May 22^nd^ 2024) using MacSyFinder (v.2.1.1; default parameters)^103^. To build HMM profiles, a PSI-BLAST of each component was performed against the NCBI non-redundant protein sequence database (accessed in February 2023, and May 2024). The PSI-blast e-value threshold was set to 1e-10 and run for three iterations, except for *Vc*SduA for which only one iteration was performed due to NCBI computational constraints. Sequences were aligned using MAFFT (v.7.508; --maxiterate 1000 –localpair parameters for a higher accuracy alignment) and HMM profiles built using the hmmbuild function of the HMMER suite (v.3.3.2; default parameters)^101,104^. To evaluate the distribution of each system in the RefSeq database used, genera represented with more than 500 genomes were extracted, and the NCBI common tree tool used to create an order-level phylogenetic tree showing the most represented clades, with the presence/absence of each system mapped. For the full list of identified hits see Supplementary Tables 3-5.

### Isolation and characterisation of Bas24 escaper phages that overcome GrwAB

Candidate escaper phages of Bas24, which appeared as spontaneous large plaques in the lowest dilution of the small-drop plaque assay on strains producing GrwAB, were isolated and propagated using a protocol modified from^105^. Single plaques of each candidate were resuspended in 90µL phage buffer (50mM Tris-HCL pH 7.5, 100 mM MgCl_2_, 10mM NaCl), added to 1mL exponentially growing *E. coli* MG1655Δ*araCBAD* producing GrwAB in a 14mL test-tube and cultured at 37°C, 180rpm for 3h, at which point a further 2mL of exponentially growing culture was added and incubated for 3h, before phages were purified exactly as described above. As a control, a single plaque from the no system control plate was picked and propagated in the absence of GrwAB production. Genomic DNA was prepared using the Norgen Phage DNA Isolation Kit (Product #46850; Norgen, Canada) according to the manufacturer’s instructions. Whole genome sequencing was done using the Small, 200 Mbp Illumina short read service on the Illumina NextSeq2000 platform (SeqCoast Genomics, U.S.A). Genomes were assembled using Unicycler (v.0.5.0)^106^, using short reads only as input with the remaining parameters set to default, and assemblies rotated to match the Bas24 reference genome^107^. Genomes were aligned using MAFFT (v.7.508) and single nucleotide polymorphisms called using snp-sites (v.2.5.1)^101,108^.

### Search for DpdA-encoding vibriophages

To identify previously described vibriophages with modified genomes, phage sequences were downloaded from the NCBI virus database, with Vibrionaceae (taxid: 641) specified as the host. Using the resulting 1,700 nucleotide assemblies and the NCBI tool suite (v.2.14.1) to translate the sequences, a local database containing 117,034 protein sequences was generated (makeblastdb). This database was then searched for homologues of DpdA from phage Bas24 using Blastp (e-value 1e-10, rest of parameters to default). The resulting data are provided in Supplementary Table 6.

### Genome sequencing of Swiss WASA-1 strain N19-2759

*V. cholerae* strain N19-2759 is a non-O1/non-O139 non-toxigenic travel-associated clinical isolate isolated from a patient in Switzerland in 2019 (Illumina-based sequencing accession number GCA_032819675.1)^78^. To obtain an assembled complete genome of strain N19-2759, genomic DNA was prepared as previously described^109^, sequenced using ONT-based sequencing, and assembled using the software flye^110^ [v. 2.9.3] (Microsynth, Switzerland).

### Data availability

The genome assemblies of *V. cholerae* strain N19-2759 have been deposited in the NCBI GenBank database under accession number [GCA_046097525.1](https://www.ncbi.nlm.nih.gov/datasets/genome/GCF_046097525.1/). The raw reads are available from the Sequence Read Archive (SRA) under submission number SRX26909066. All other data are available in the main text or the supplementary materials.

## Acknowledgments

We thank all the members of the Blokesch laboratory for constructive feedback throughout the project. We also gratefully acknowledge K. D. Seed (University of California, Berkeley), A. Camilli (Tufts University School of Medicine, Boston), W.-L. Ng (Tufts University School of Medicine, Boston) and A. Ali (University of Florida, Gainesville) for sharing ICP phages, and A. Harms (ETH, Zurich) for sharing the BASEL collection. We thank the Félix d’Hérelle Reference Center for Bacterial Viruses from the Université Laval (Québec, Canada) for providing phage X29 (HER66). Finally, we acknowledge Roger Stephan and Michael Biggel (University of Zurich) for sharing *V. cholerae* isolates from Switzerland and for valuable scientific discussions.

## Author contributions

Conceptualization: DWA, MJ and MB

Methodology: DWA, MJ, AL, and MB

Investigation: DWA, MJ, AL, SS, LR, LB and MB

Visualization: DWA

Funding acquisition: MB

Supervision: DWA and MB

Writing – original draft: DWA

Writing – review & editing: DWA, MJ and MB

Writing – revision: DWA and MB

## Supplementary information

**Extended Data Fig. 1.**
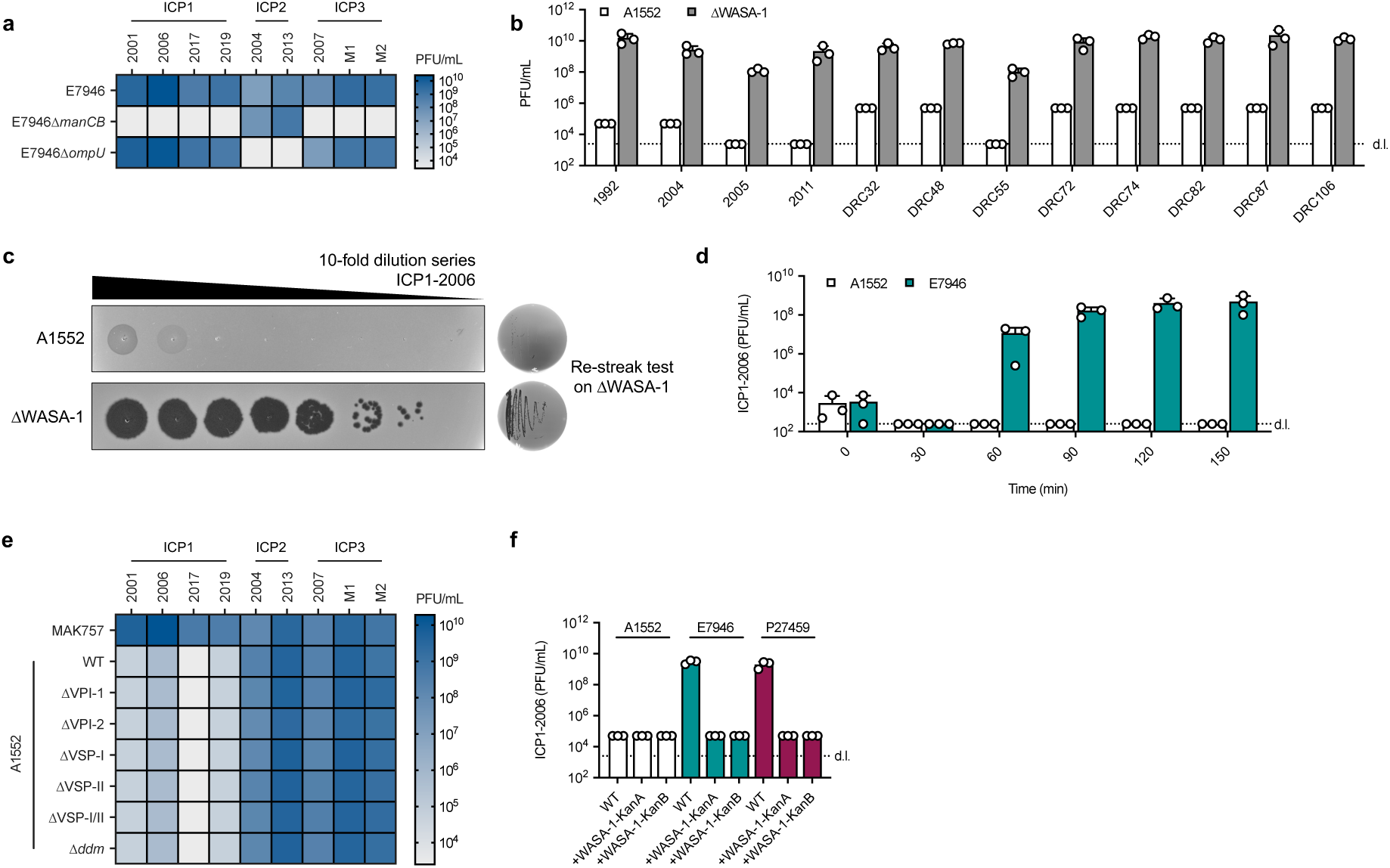
Control experiments showing role of WASA-1 prophage in ICP1 defence. (**a**) Heat-map showing mean plaque-forming units (PFU/mL) of diverse ICP1-3 isolates determined by plaque assays on E7946 derivatives that lack either the O1 antigen receptor for ICP1 and ICP3 (E7946Δ*manCB*) or the OmpU receptor for ICP2 (E7946Δ*ompU*). (**b**) Plaque assay showing PFU/mL of various ICP1 isolates on A1552 in the presence and absence of WASA-1. (**c**) Re-streak tests evaluating the propagation of viable ICP1 phage progeny were conducted by taking material from the highest concentration spot of 10-fold serial dilution plaque assays done in the presence (A1552) and absence (ΔWASA-1) of WASA-1 (left) and re-streaking on fresh lawns of the susceptible ΔWASA-1 strain (right). (**d**) Replication assay showing change in PFU/mL over time following the infection of A1552 and E7946 cultures with *c.a.* 10^3^-10^4^ PFU of ICP1-2006. (**e**) Heat-map showing mean plaque-forming units (PFU/mL) of diverse ICP1-3 isolates determined by plaque assays on A1552 derivatives that lack the indicated genomic islands, as compared to the pre-7^th^ pandemic control strain MAK757. (**f**) Plaque assay showing the effects of introducing WASA-1, using versions with a kanamycin resistance cassette inserted at two different positions, into the non-WASA lineage strains E7946 and P27459 on ICP1-2006 PFU/mL, as compared to the equivalent A1552 control strains. Heat-maps and bar charts show the mean from 3 independent repeats. Error bars show s.d. and d.l. = detection limit.

**Extended Data Fig. 2.**
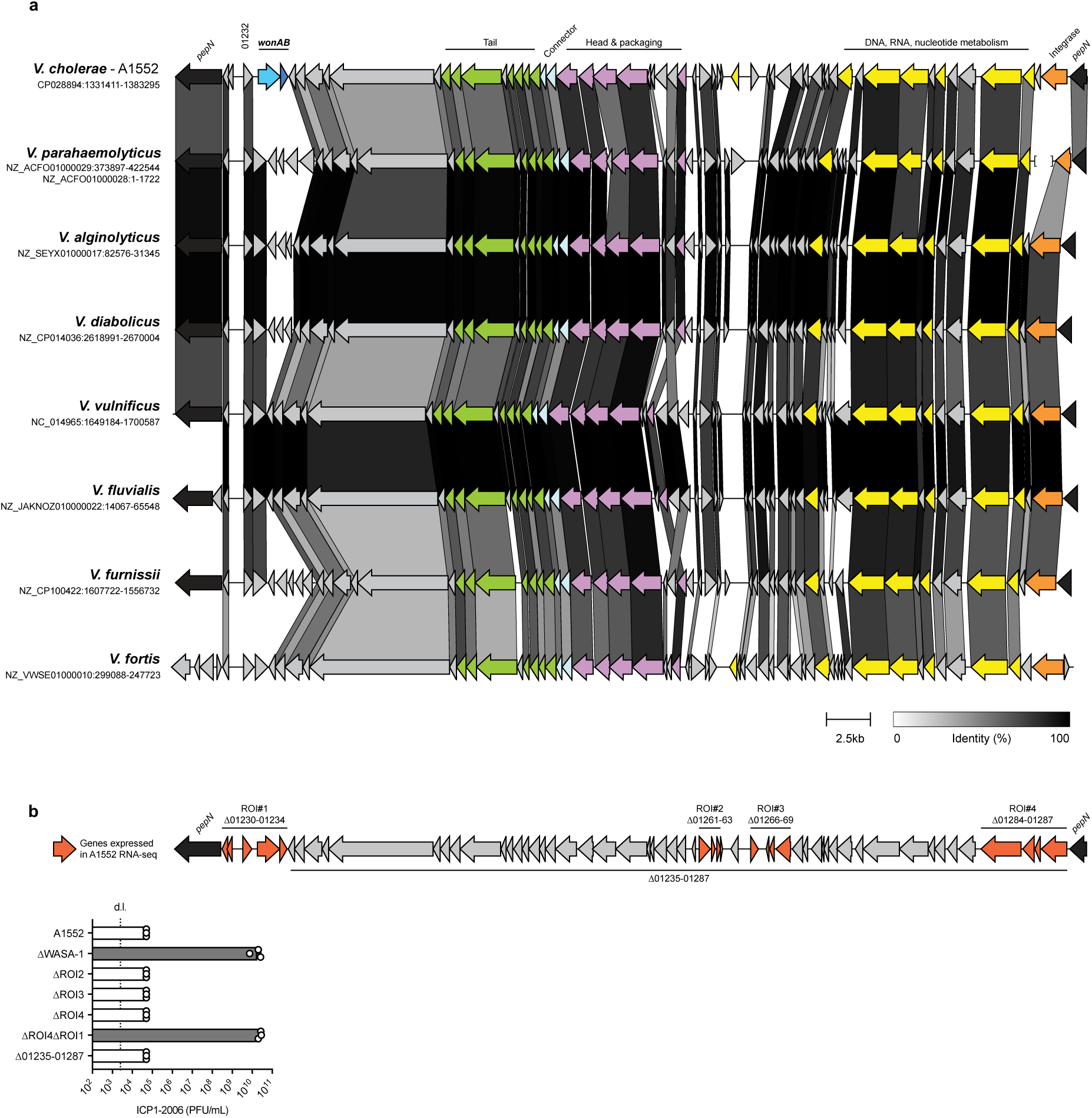
Conservation of the WASA-1 genome and identification of regions of interest. (**a**) Schematic comparing the genome organisation of WASA-1 from *V. cholerae* strain A1552 with a representative set of examples of the WASA-1 detected by BLAST in diverse *Vibrio spp*. The schematic was built using the clinker pipeline and the connections between genomes are coloured according to protein identity (above the minimum default threshold of 30%). Genes with predicted functions in prophage biology are coloured according to function, as indicated. For details of WASA-1 identification and a full list of identified sequences, see methods and Supplementary Table 2. (**b**) Schematic of *V. cholerae* strain A1552 WASA-1 highlighting the genes that were highly expressed under standard growth conditions in RNA-seq results, and which were used to define the indicated regions of interest (ROI) 1-4. Note that ROI1 could only be deleted in the ΔROI4 background. The bar chart shows the effects of deleting each ROI on ICP1-2006 PFU/mL, as compared to WT and ΔWASA-1 control strains. Bars show the mean + s.d. from 3 independent repeats. d.l. = detection limit.

**Extended Data Fig. 3.**
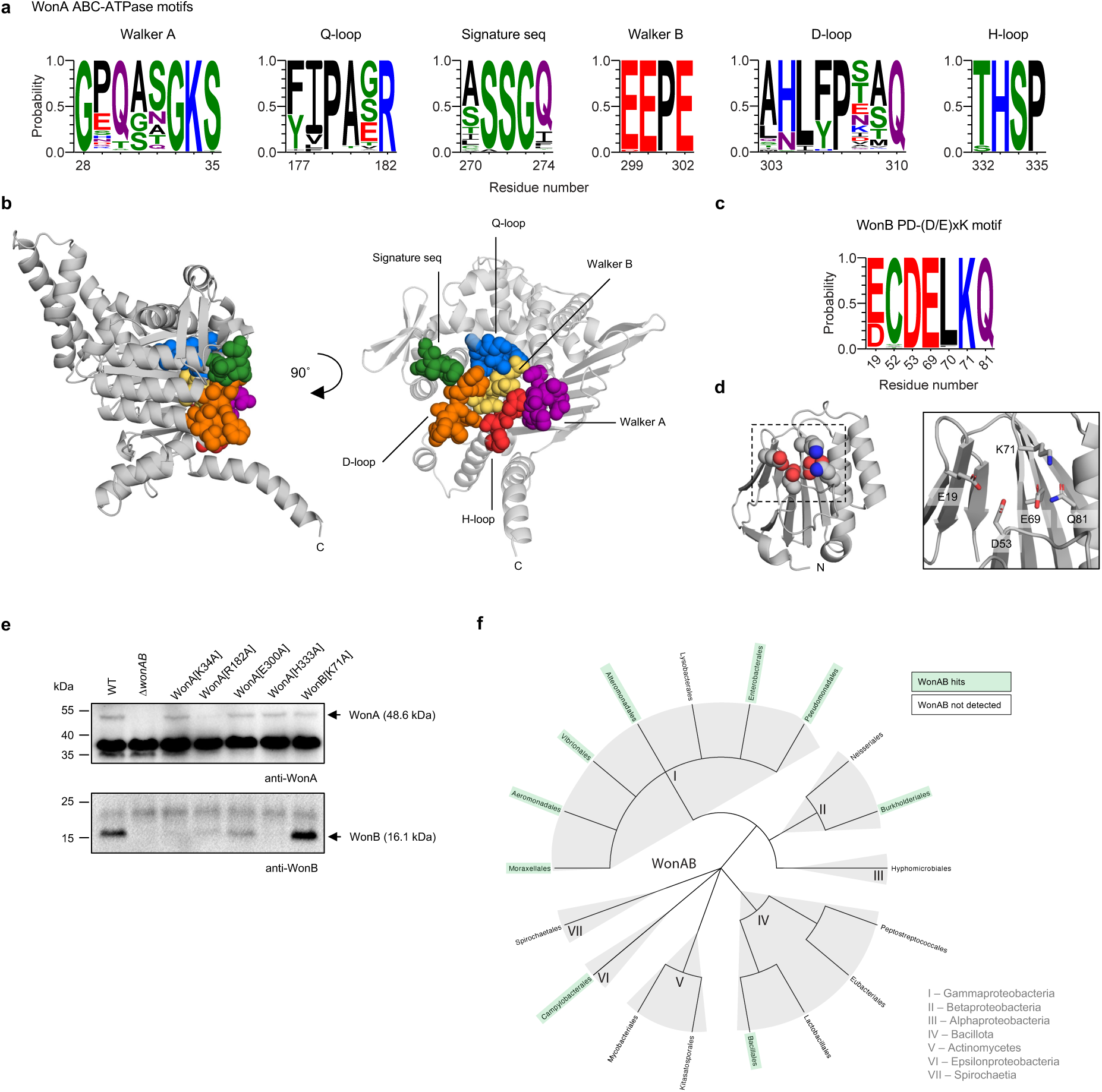
Identification of conserved motifs and distribution of WonAB. (**a**) Sequence logos showing conservation of the identified ABC-ATPase motifs in the 198 WonA hits detected using MacSyFinder v.2.1.1, as compared to the equivalent residue number in *V. cholerae* A1552 WonA. Amino acids in logos are coloured according to chemical properties: polar (G, S, T, Y, C), green; neutral (Q, N), purple; basic (K, R, H), blue; acidic (D, E), red; and hydrophobic (A, V, L, I, P, W, F, M), black. (**b**) Location of the conserved ABC-ATPase motifs shown in (**a**) at the dimer interface of the WonA dimer structural prediction. (**c**) Sequence logo showing the conservation of the identified PD-(D/E)xK nuclease motif in the 198 WonB hits detected using MacSyFinder v.2.1.1, compared to the equivalent residue number in *V. cholerae* A1552 WonB. (**d**) Location of the identified PD-(D/E)xK nuclease residues in the WonB structural prediction. (**e**) Western blot showing the protein levels of WonA and WonB, natively expressed in strain A1552 (WT) and in derivatives encoding the indicated site-directed variants. (**f**) Distribution of WonAB hits detected using MacSyFinder v.2.1.1. The tree shows the order-level phylogeny of genera in the RefSeq database with more than 500 genomes (see methods). For the full list of 198 WonAB hits see Supplementary Table 3.

**Extended Data Fig. 4.**
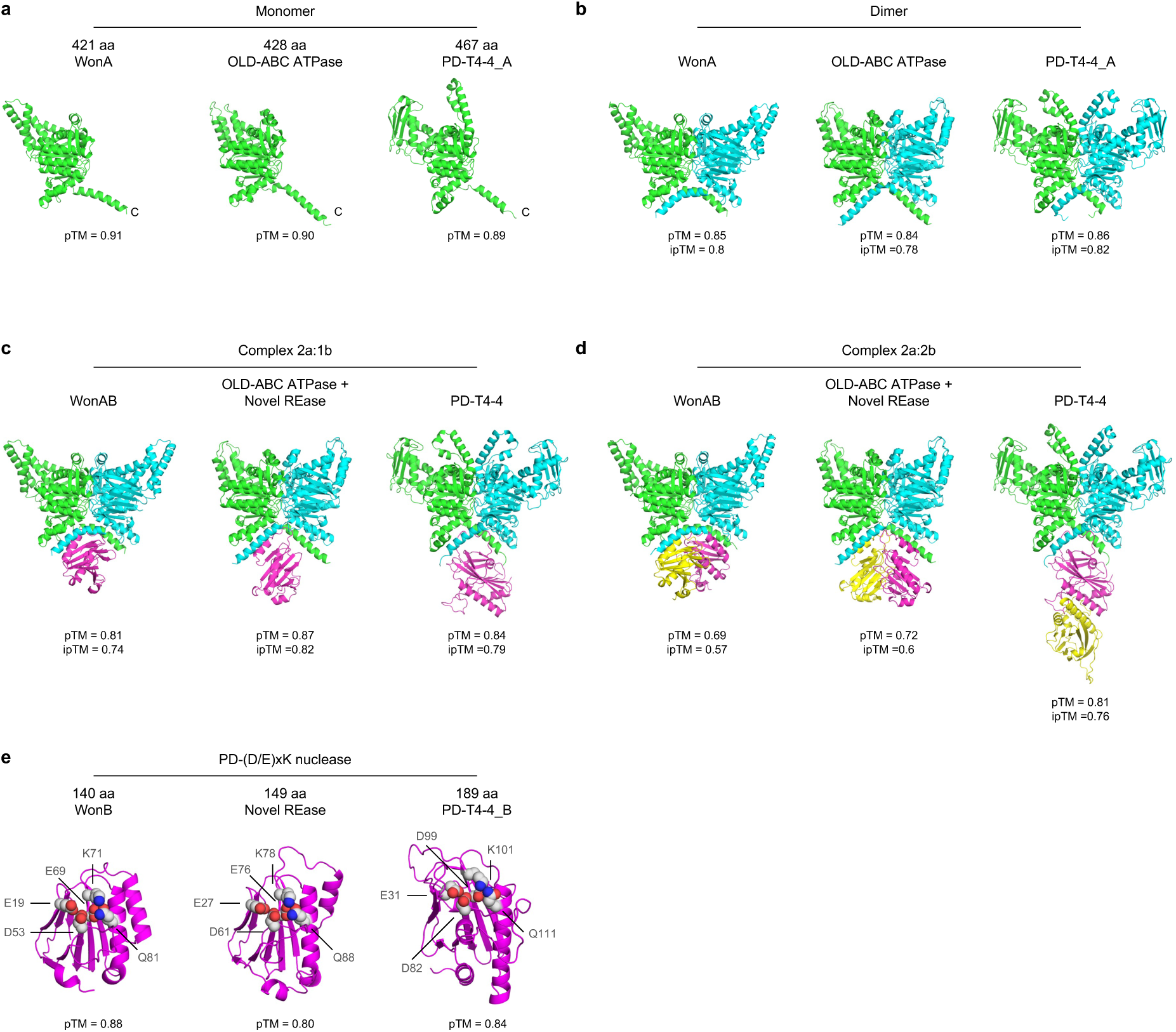
Summary of structural modelling of WonAB and related systems. (**a-e**) Cartoon representations coloured by chain showing the AlphaFold3 predicted structures for WonAB, and the related OLD-ABC ATPase + Novel REase and PD-T4-4 systems. The ATPase components WonA, OLD-ABC ATPase and PD-T4-4_A were modelled as a monomer (**a**) and a dimer (**b**), and further modelled as a complex with either one (**c**) or two (**d**) copies of the nuclease components WonB, Novel REase and PD-T4-4_B. (**e**) Nuclease components modelled alone, highlighting the location of the predicted PD-(D/E)xK nuclease residues. The predicted template modelling (pTM) score, and where appropriate, the interface predicted template modelling (ipTM) score, are shown alongside each model. Protein sequences were obtained from NCBI: *V. cholerae* A1552 WonA (AWB73975.1) and WonB (AWB73976.1); *Anaerovibrio lipolyticus* DSM 3074 OLD-ABC ATPase (SHI83489.1) + Novel REase (SHI83462.1); *E. coli* MOD1-ECOR58 PD-T4-4_A (RCO57999.1) and PD-T4-4_B (RCO57988.1).

**Extended Data Fig. 5.**
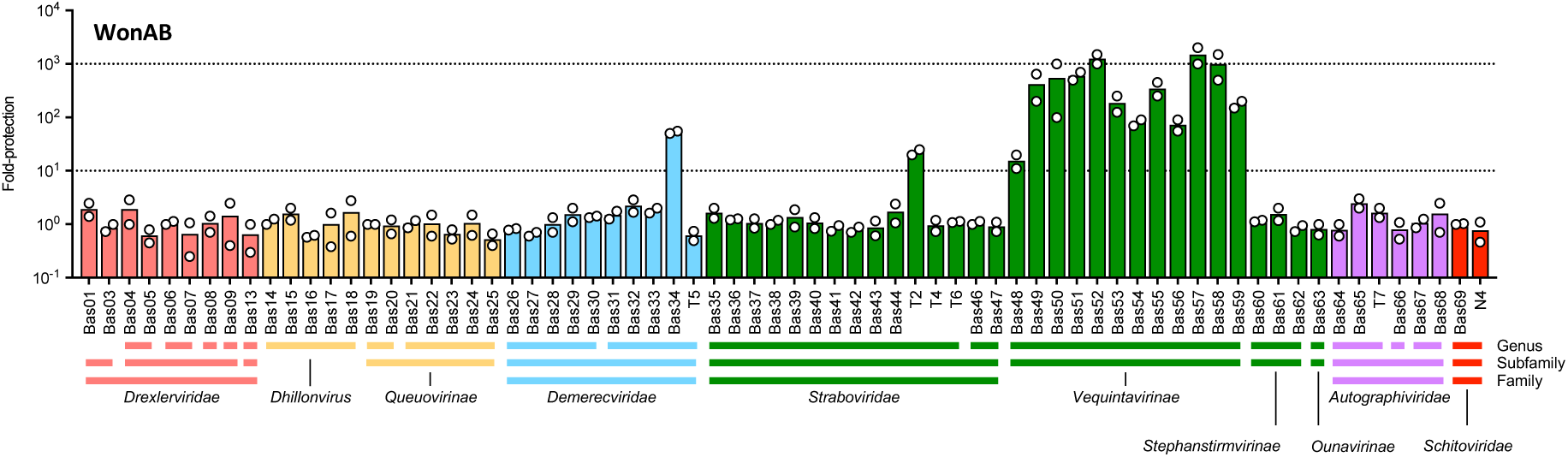
Anti-phage activity of WonAB in *E. coli*. Fold-protection against *E. coli* phages of the BASEL collection conferred by the production of WonAB in *E. coli* MG1655Δ*araCBAD*, as compared to a negative ‘no system’ control. The system was expressed from a chromosomally integrated transposon carrying the arabinose-inducible *P*_BAD_-promoter, induced by the addition of 0.2% arabinose. Bar chart shows the mean of two independent experiments.

**Extended Data Fig. 6.**
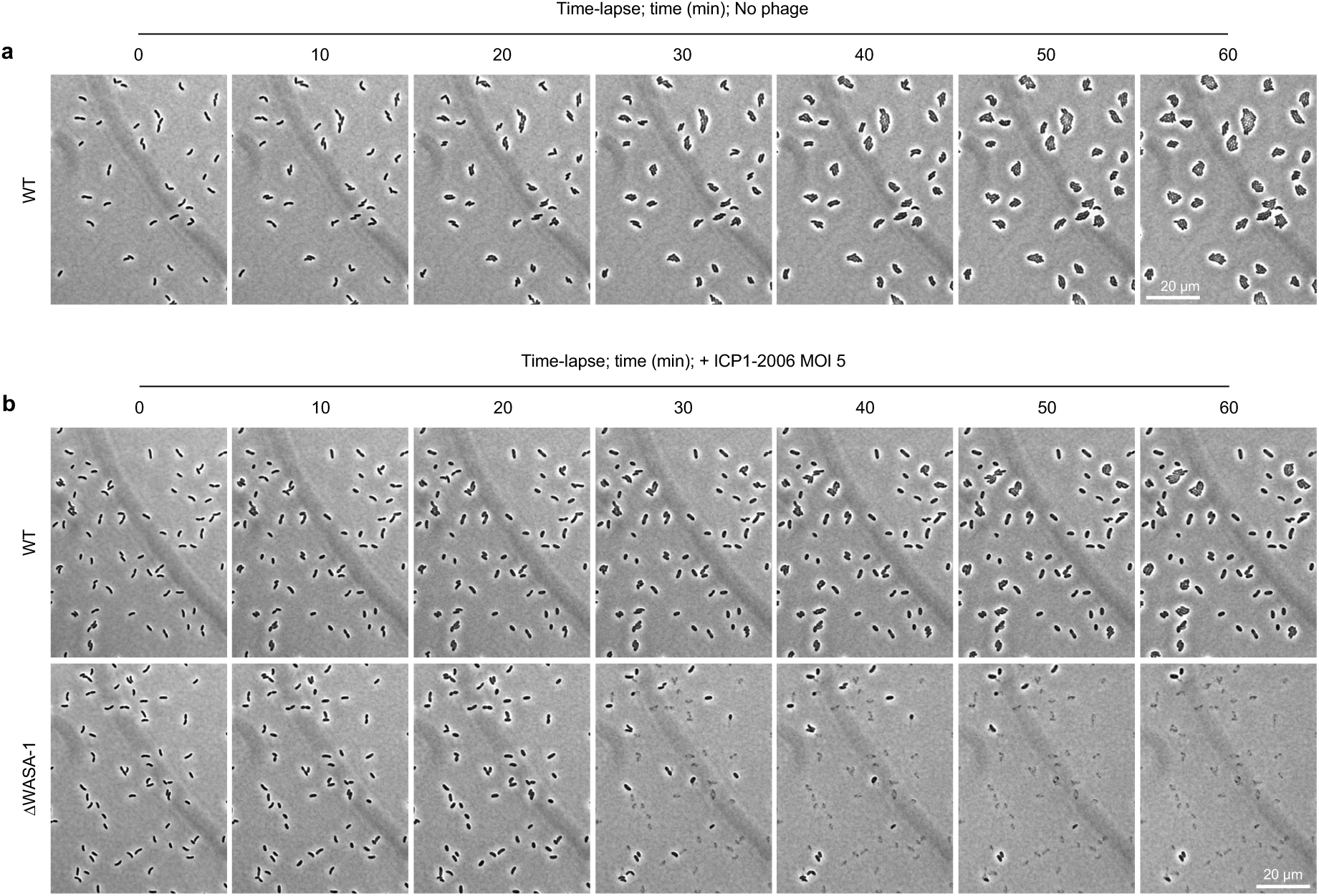
Time-lapse microscopy of ICP1 infection. (**a**) Control experiment showing time-lapse microscopy of exponentially growing cells of *V. cholerae* A1552 (WT) in the absence of phage infection. (**b**) Time-lapse microscopy comparing exponentially growing cells of *V. cholerae* WT and ΔWASA-1 strains after infection with ICP1-2006 at MOI 5. The panels depict the full-frame versions of the examples presented in Fig. 2b. All images are representative of the results of three independent experiments. Scale bars = 20 µm.

**Extended Data Fig. 7.**
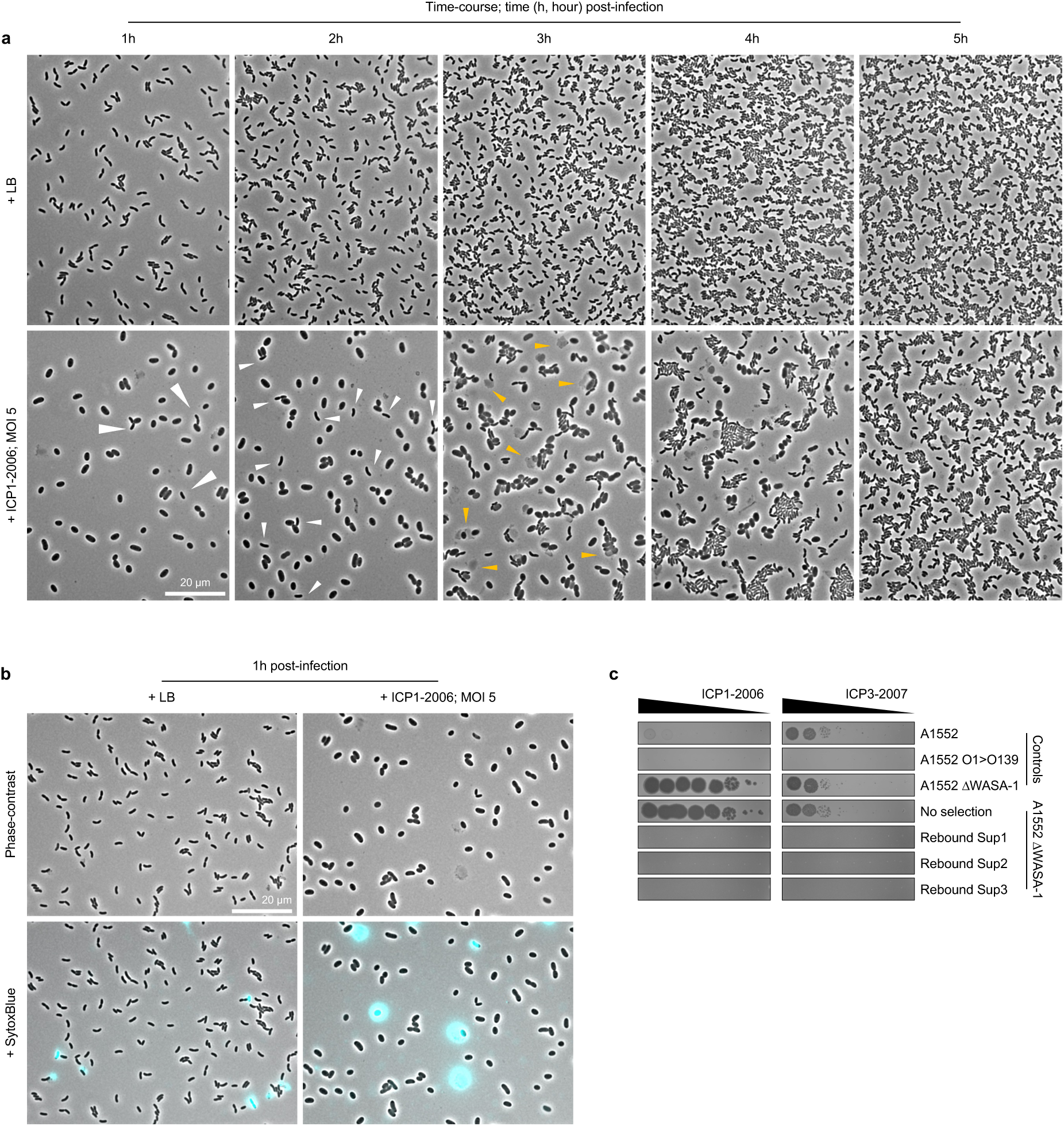
Imaging the fate of ICP1 infected cells. (**a**) Time-course microscopy snapshots comparing the growth and cell morphology of *V. cholerae* A1552 cultures at 1, 2, 3, 4 and 5 hours after infection with ICP1-2006 at MOI 5, as compared to an uninfected control culture (+ LB). Note how the ICP1-infected culture initially contains only a few cells with normal morphology (white arrowheads), and that these cells increase in number as the culture rebounds, while the swollen ICP1-infected cells that are initially dominant persist for several hours before undergoing lysis (yellow arrowheads). (**b**) SytoxBlue-staining of *V. cholerae* A1552 cultures 1 hour after infection with ICP1-2006 at MOI 5, as compared to an uninfected control culture (+ LB). (**c**) Plaque assays testing the ability of ICP1-2006 and ICP3-2007 (which both require the O1 antigen receptor) to form plaques on cultures of A1552ΔWASA-1 that rebounded following ICP1-infection, as compared to a culture of the same strain that was grown without selection, and the set of indicated control strains. Note how neither ICP1-2006 nor ICP3-2007 are able to form plaques on the rebounded cultures, consistent with recovery of the culture being driven by the growth of spontaneous O1-antigen mutant cells. All images are representative of the results of three independent experiments. Scale bars = 20 µm.

**Extended Data Fig. 8.**
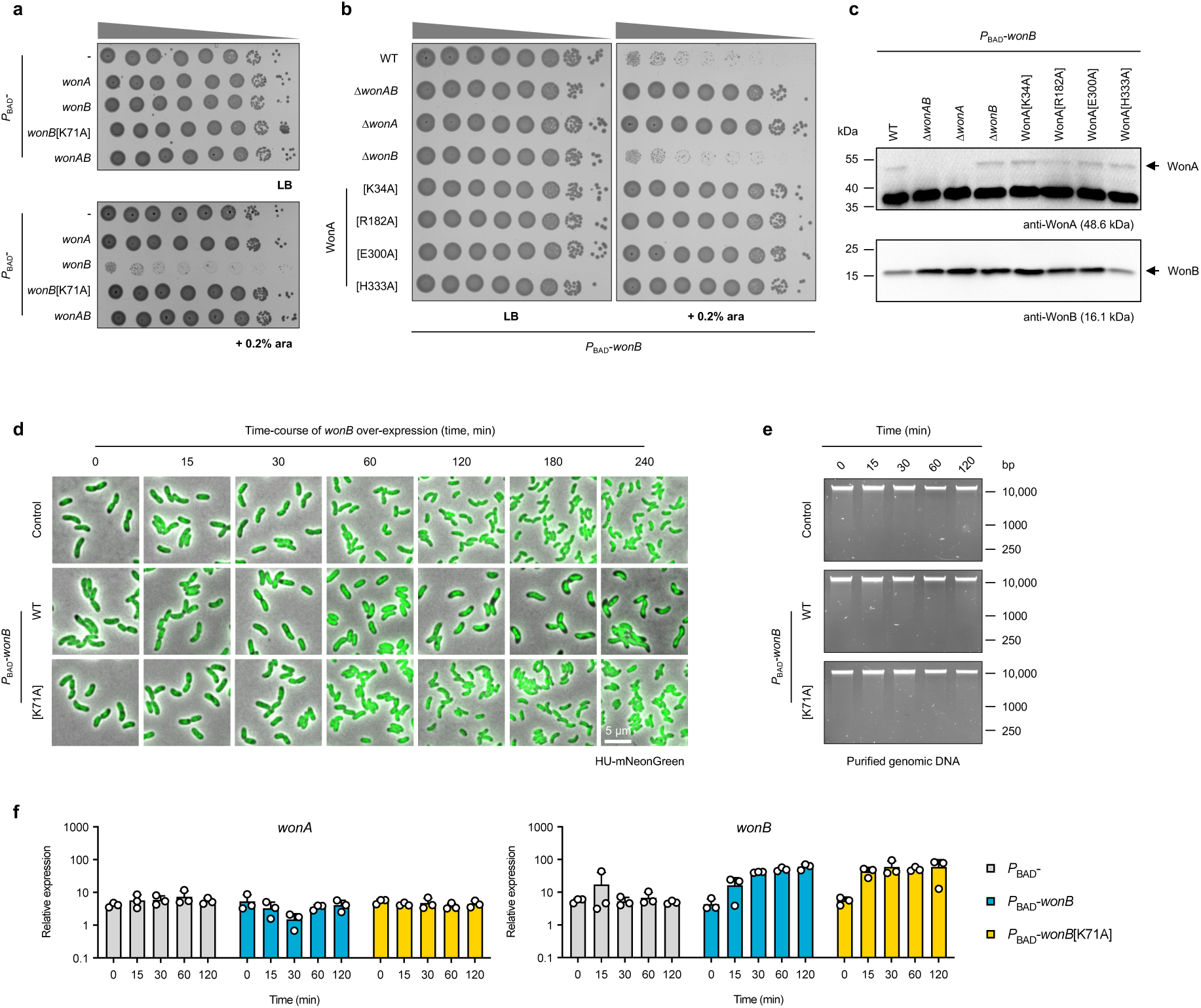
Control experiments showing details of toxicity during WonB overproduction. (**a**) Toxicity assay evaluating the growth of 10-fold serial dilutions of cultures of *V. cholerae* strain A1552 derivatives with a chromosomally integrated transposon carrying the indicated genes under the control of the arabinose-inducible *P*_BAD_-promoter, in the absence (LB) and presence of induction (+ 0.2% ara), as compared to a negative control strain. (**b**) Toxicity assay evaluating the growth of 10-fold serial dilutions of *V. cholerae* strain A1552 and the indicated derivatives encoding site-directed variants of WonA, compared to the indicated deletion control strains, in the absence (LB) and presence (+ 0.2% ara) of WonB overproduction. Note that the control strains are also shown separately in Fig. 3a. (**c**) Western blots showing the protein levels of WonA and WonB in the strains used for the toxicity assay shown in (**b**). Samples were taken from exponentially growing cultures, 15 minutes after induction of *P*_BAD_-*wonB*. (**d**) Time-course microscopy snapshots showing the effect of WonB overproduction on cell morphology and cellular DNA content, as monitored by a HupA-mNeonGreen fusion, compared to a negative control strain and a strain overproducing the inactive WonB[K71A] variant. Scale bar = 5 µm. The panels depict the full time-series for the examples presented in Fig. 3d. (**e**-**f**) Agarose gels evaluating the integrity of genomic DNA extractions (**e**) and comparison of *wonA* and *wonB* transcript levels as determined by qRT-PCR (**f**), prepared from cultures over time upon WonB overproduction following induction in exponentially growing cells at time 0, as compared to a negative control strain and a strain overproducing the inactive WonB[K71A] variant. All data are representative of the results of three independent repats. Bar charts show the mean + s.d.

**Extended Data Fig. 9.**
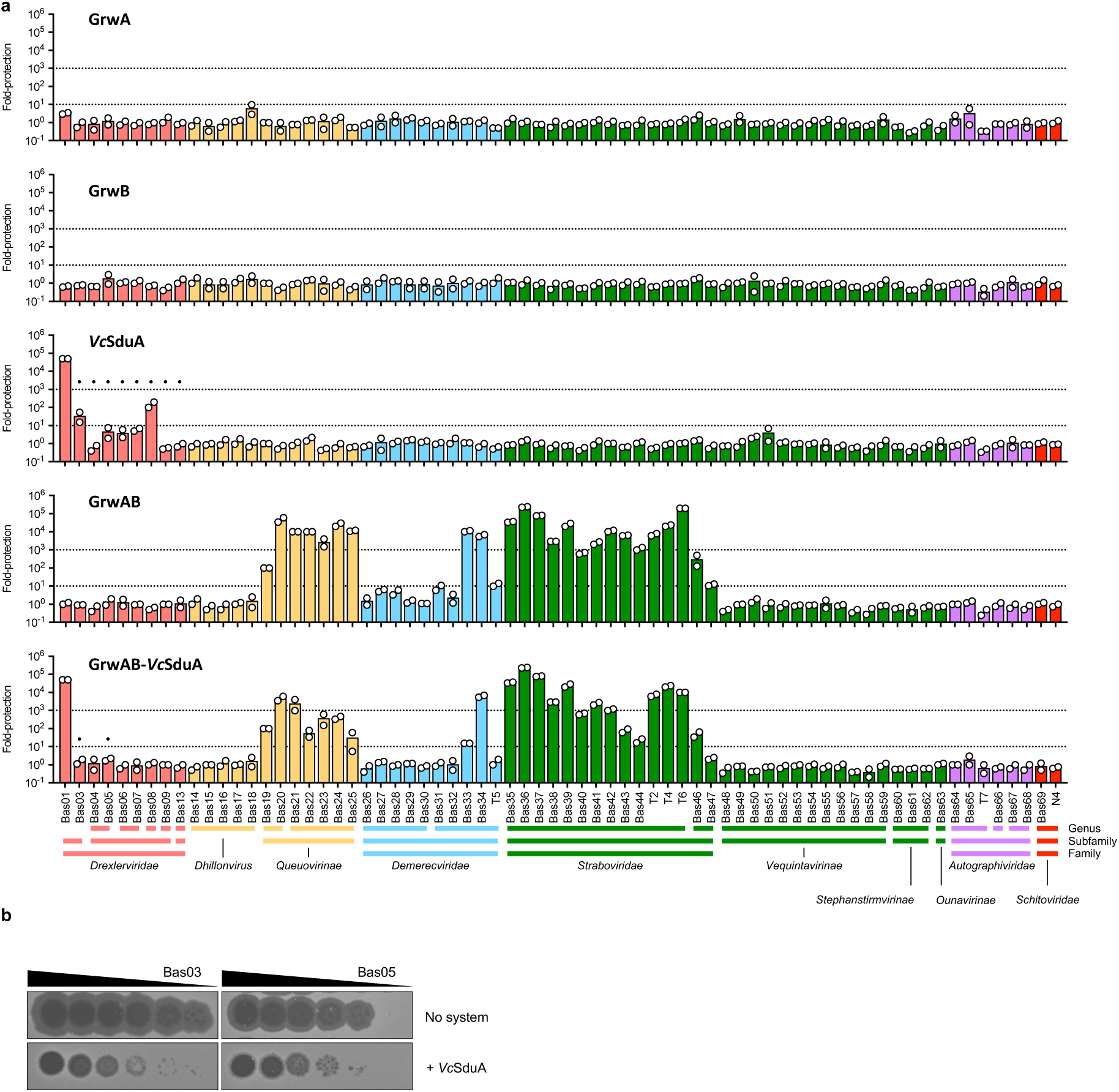
Anti-phage activity of GrwAB and *Vc*SduA in *E. coli*. **(a)** Fold-protection against *E. coli* phages of the BASEL collection conferred by the production of either GrwA, GrwB, *Vc*SduA, GrwAB or GrwAB-*Vc*SduA in *E. coli* MG1655Δ*araCBAD*, as compared to a negative ‘no system’ control. Genes were expressed from a chromosomally integrated transposon carrying the arabinose-inducible *P*_BAD_-promoter, induced by the addition of 0.2% arabinose. Bar charts show the mean of two independent experiments. Bars with a black dot indicate phages exhibiting altered plaque morphology. (**b**) Representative examples of the altered plaque morphology phenotype observed for Bas03 and Bas05 in the presence of *Vc*SduA production.

**Extended Data Fig. 10.**
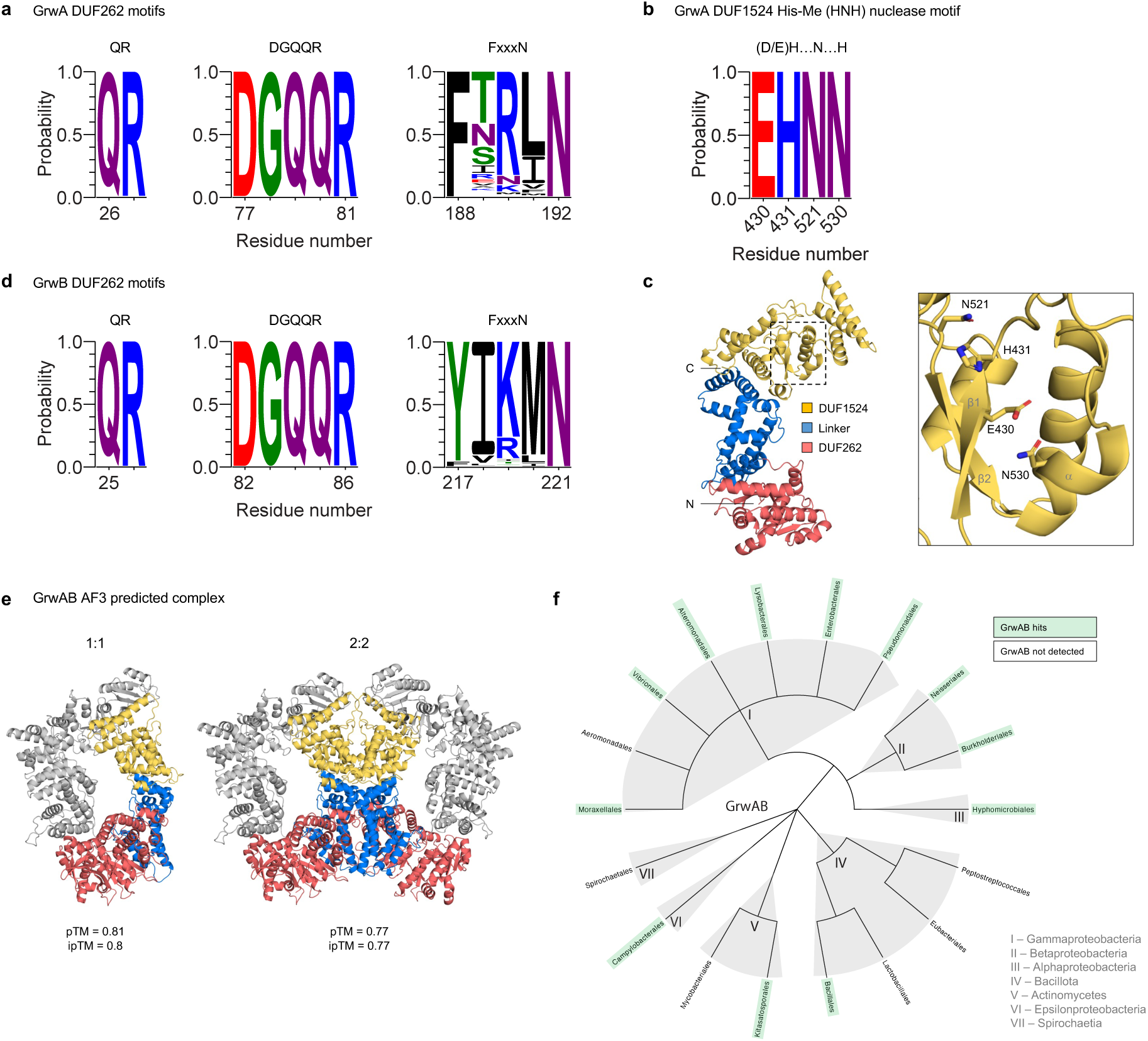
Identification of conserved motifs and distribution of GrwAB. (**a**, **d**) Sequence logos showing conservation of the identified QR, DGQQR and FxxxN motifs in the DUF262 domains of the 471 GrwA (**a**) and 471 GrwB (**d**) hits detected using MacSyFinder v.2.1.1, as compared to the equivalent residue number in *V. cholerae* A1552 GrwA and GrwB. Amino acids in logos are coloured according to chemical properties: polar (G, S, T, Y, C), green; neutral (Q, N), purple; basic (K, R, H), blue; acidic (D, E), red; and hydrophobic (A, V, L, I, P, W, F, M), black. (**b**) Sequence logo showing the conservation of the identified His-Me (HNH) nuclease motif in the DUF1524 domain of the 471 GrwA hits detected using MacSyFinder v.2.1.1, as compared to the equivalent residue number in *V. cholerae* A1552 GrwA. (**c**) Location of the identified His-Me (HNH) nuclease motif residues in the GrwA structural prediction, highlighting the predicted ββα fold. (**e**) AlphaFold3 predicted structures of potential GrwAB complexes. Colour scheme as in (**c**). (**f**) Distribution of GrwAB hits detected using MacSyFinder v.2.1.1. The tree shows the order-level phylogeny of genera in the RefSeq database with more than 500 genomes (see methods). For the full list of 471 GrwAB hits see Supplementary Table 4.

**Extended Data Fig. 11.**
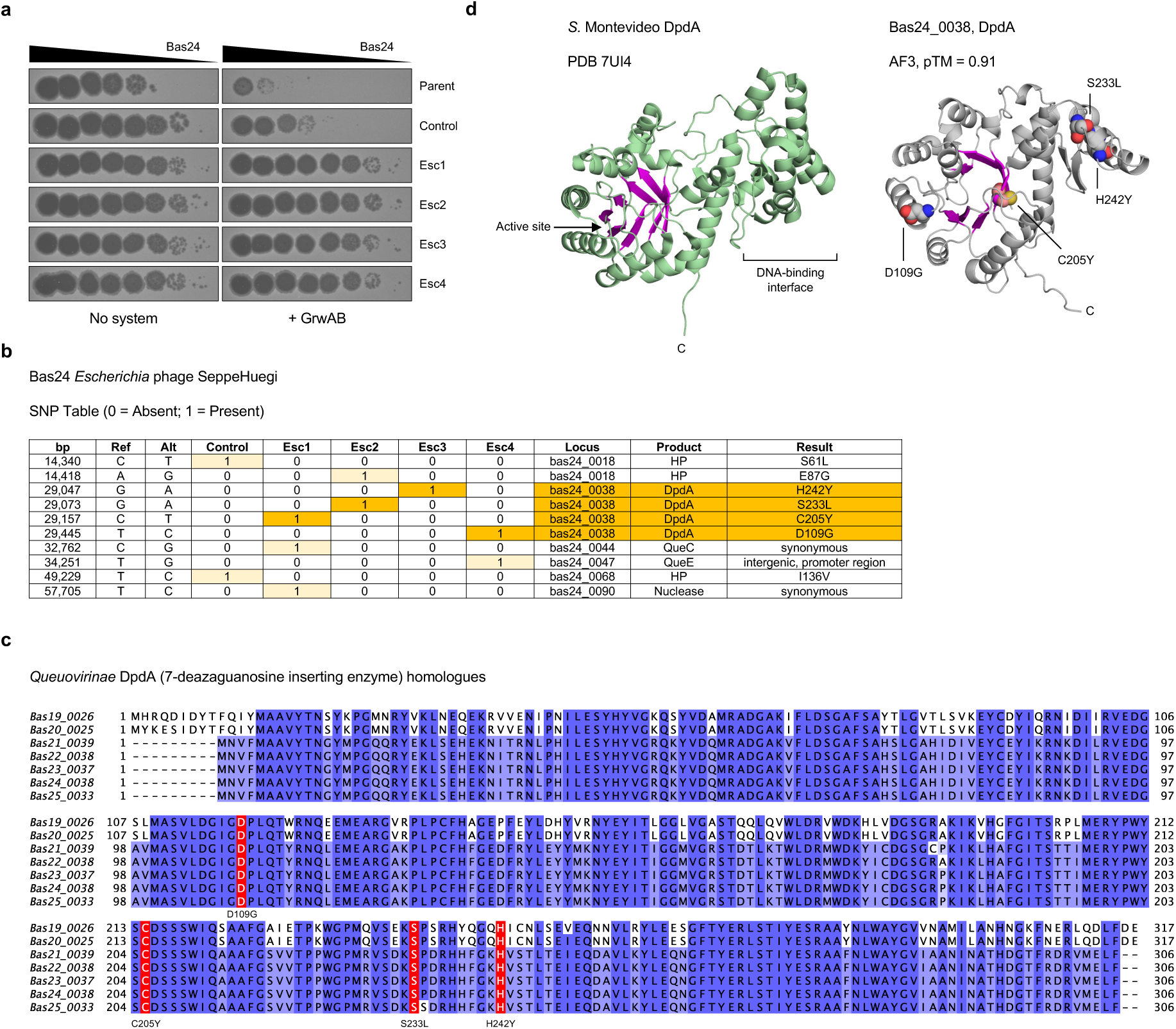
Bas24 escaper phages overcome GrwAB defence via mutations in the DNA modification pathway. (**a**) Plaque assays evaluating the ability of 10-fold serial dilutions of Bas24 escaper phages to form plaques on *E. coli* MG1655Δ*araCBAD* in the absence (No system) and presence of GrwAB (+ GrwAB), as compared to the parental Bas24 stock and a control stock propagated in the absence of GrwAB. (**b**) SNP table summarising results of whole genome sequencing for each phage compared to the published Bas24 reference sequence. Light yellow (SNP), dark yellow (SNP targeting *dpdA* locus), HP (hypothetical protein). (**c**) Multiple-sequence alignment showing conservation of *Queuovirinae* (Bas19-25) homologues of the 7-deazaguanosine inserting enzyme DpdA, with the residues found to be substituted in the Bas24 escaper phages highlighted in red. (**d**) Locations of the substituted residues shown on predicted structure of Bas24 DpdA, generated using AlphaFold3, compared to the experimentally determined crystal structure of *Salmonella enterica* serovar Montevideo DpdA (PDB 7UI4) showing the active site highlighted in magenta.

**Extended Data Fig. 12.**
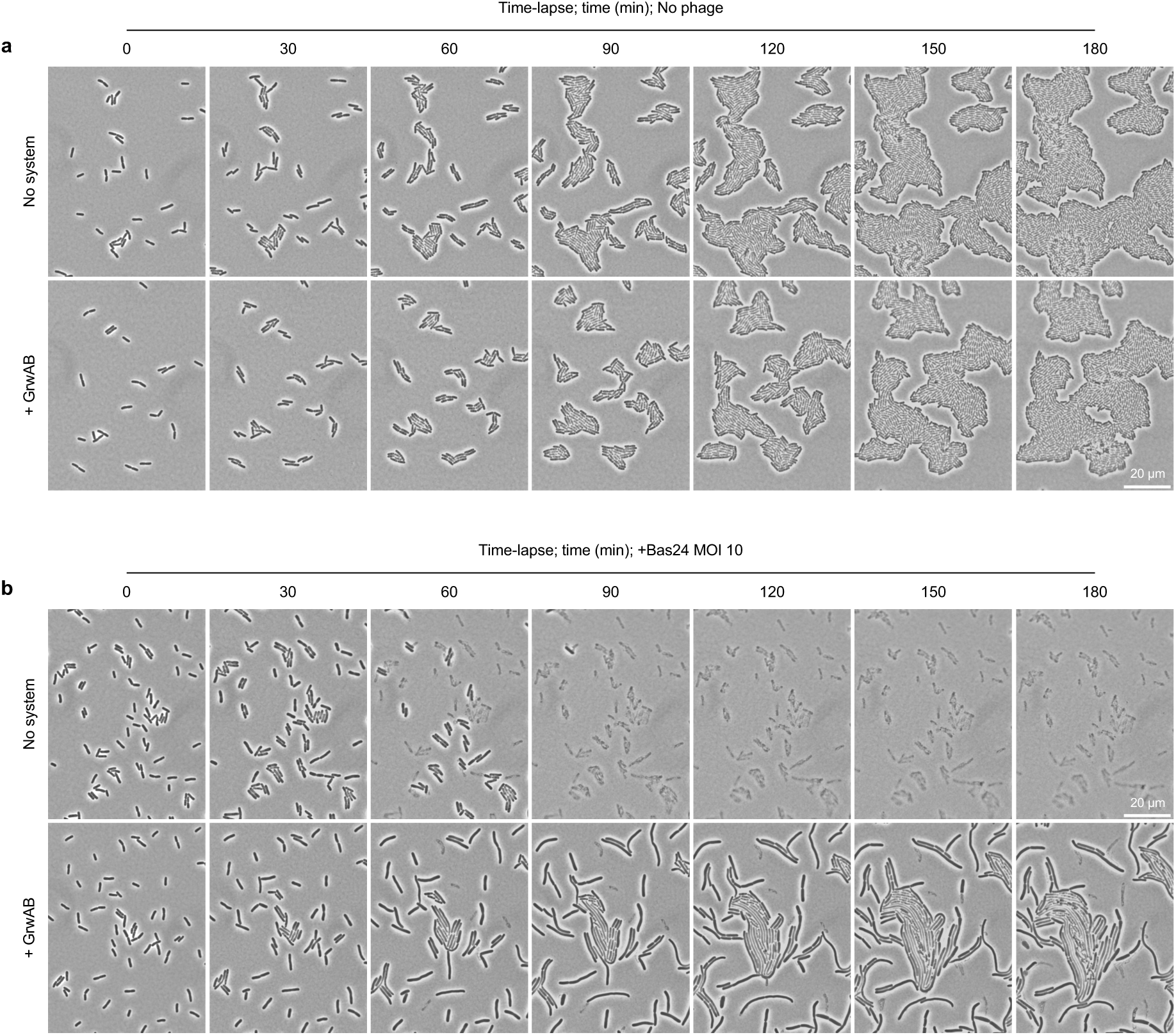
Time-lapse microscopy of Bas24 infection. (**a**-**b**) Time-lapse microscopy of exponentially growing cells of *E. coli* MG1655Δ*araCBAD* in the absence (No system) and presence (+ GrwAB) of GrwAB production, in the absence (**a**) and presence (**b**) of infection with Bas24 at MOI 10. All images are representative of the results of three independent experiments. Scale bars = 20 µm.

**Extended Data Fig. 13.**
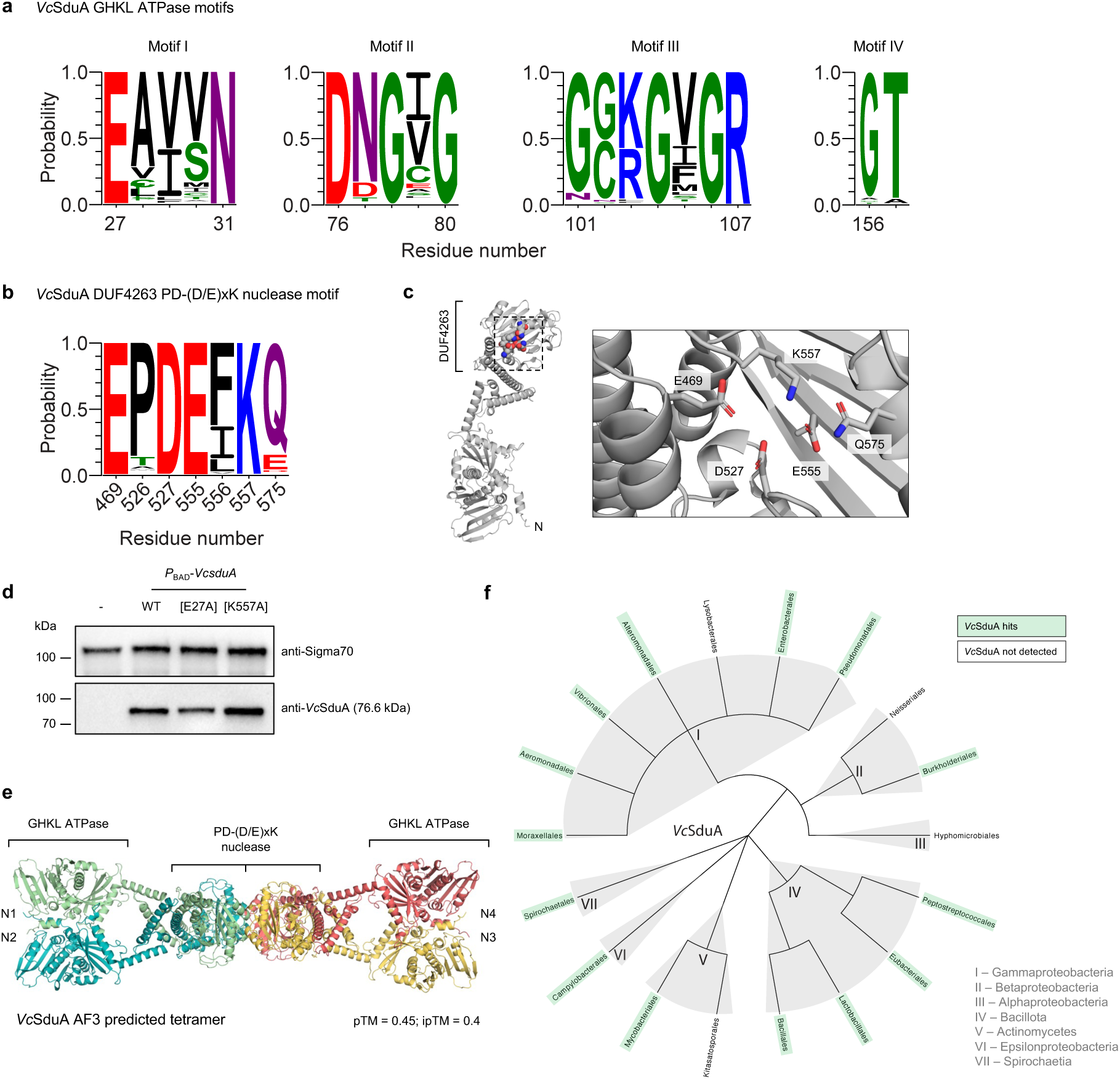
Identification of conserved motifs and distribution of *Vc*SduA. (**a**-**b**) Sequence logos showing conservation of the identified GHKL ATPase motifs I-IV in the GHKL domain (**a**) and the identified PD-(D/E)xK nuclease motif in the DUF4263 domain (**b**) of the 543 *Vc*SduA hits detected using MacSyFinder v.2.1.1, as compared to the equivalent residue number in *V. cholerae* A1552 *Vc*SduA. Amino acids in logos are coloured according to chemical properties: polar (G, S, T, Y, C), green; neutral (Q, N), purple; basic (K, R, H), blue; acidic (D, E), red; and hydrophobic (A, V, L, I, P, W, F, M), black. (**c**) Location of the identified PD-(D/E)xK nuclease residues in the *Vc*SduA structural prediction. (**d**) Western blot showing the protein levels of *Vc*SduA, expressed in *E. coli* MG1655Δ*araCBAD* from a chromosomally integrated transposon carrying the arabinose-inducible *P*_BAD_-promoter, induced by the addition of 0.2% arabinose, as compared to derivatives encoding the indicated site-directed variants. Sigma70 was used as a loading control. (**e**) AlphaFold3 predicted structure of potential *Vc*SduA tetramer. (**f**) Distribution of *Vc*SduA hits detected using MacSyFinder v.2.1.1. The tree shows the order-level phylogeny of genera in the RefSeq database with more than 500 genomes (see methods). For the full list of 543 *Vc*SduA hits see Supplementary Table 5.

### Supplementary Tables

**Table 1.** Summary of defence systems identified by DefenseFinder and PADLOC.

**Table 2.** Summary of WASA-1 hits detected by BLAST.

**Table 3.** Summary of matching hits for the *Vibrio cholerae* WonAB and *Anaerovibrio lipolyticus* OLD-ABC ATPase + Novel REase models, detected by MacSyFinder v.2.1.1.

**Table 4.** Summary of matching hits for the *Vibrio cholerae* GrwAB model detected by MacSyFinder v.2.1.1.

**Table 5.** Summary of matching hits for the *Vibrio cholerae Vc*SduA model detected by MacSyFinder v.2.1.1.

**Table 6.** Summary of DdpA-encoding vibriophages detected by BLAST.

**Table 7.** Bacterial strains, plasmids, bacteriophages and defence systems used in this work.

### Supplementary Movies

**Movie 1**. Time-lapse microscopy of ICP1 infection.

Time-lapse microscopy comparing exponentially growing cells of *V. cholerae* A1552 WT and ΔWASA-1 strains after infection with ICP1-2006 at MOI 5. Cells were grown at 37°C on LB agarose and imaged automatically at 1 minute intervals for 60 minutes. Bar = 20 µm. Playback = 5 fps.

**Movie 2**. Time-lapse microscopy of Bas24 infection.

Time-lapse microscopy comparing exponentially growing cells of *E. coli* MG1655Δ*araCBAD* in the absence (No system) and presence of GrwAB (+ GrwAB) production, after infection with Bas24 at MOI 10. Cells were grown at 37°C on LB agarose + 5mM CaCl_2_, 20mM MgSO_4_, 0.2% arabinose and imaged automatically at 1 minute intervals for 180 minutes. Bar = 20 µm. Playback = 5 fps.

### Supplementary Figures

**Supplementary Fig. 1. Confidence metrics for AlphaFold3 models.**

AlphaFold3 models of (**a**) WonAB and related systems (**b**) GrwAB, (**c**) *Vc*SduA and (**d**) DpdA are shown below predicted aligned error (PAE) plots and are coloured according to per residue predicted local distance difference test (pLDDT) score. The predicted template modelling (pTM) score, and where appropriate the interface predicted template modelling (ipTM) score, are shown below each model.

