## Supplementary figures and images for "Diverse phage defence systems define West African South American pandemic *Vibrio cholerae*"

### Supplementary Figure 1

**Supplementary Fig. 1**

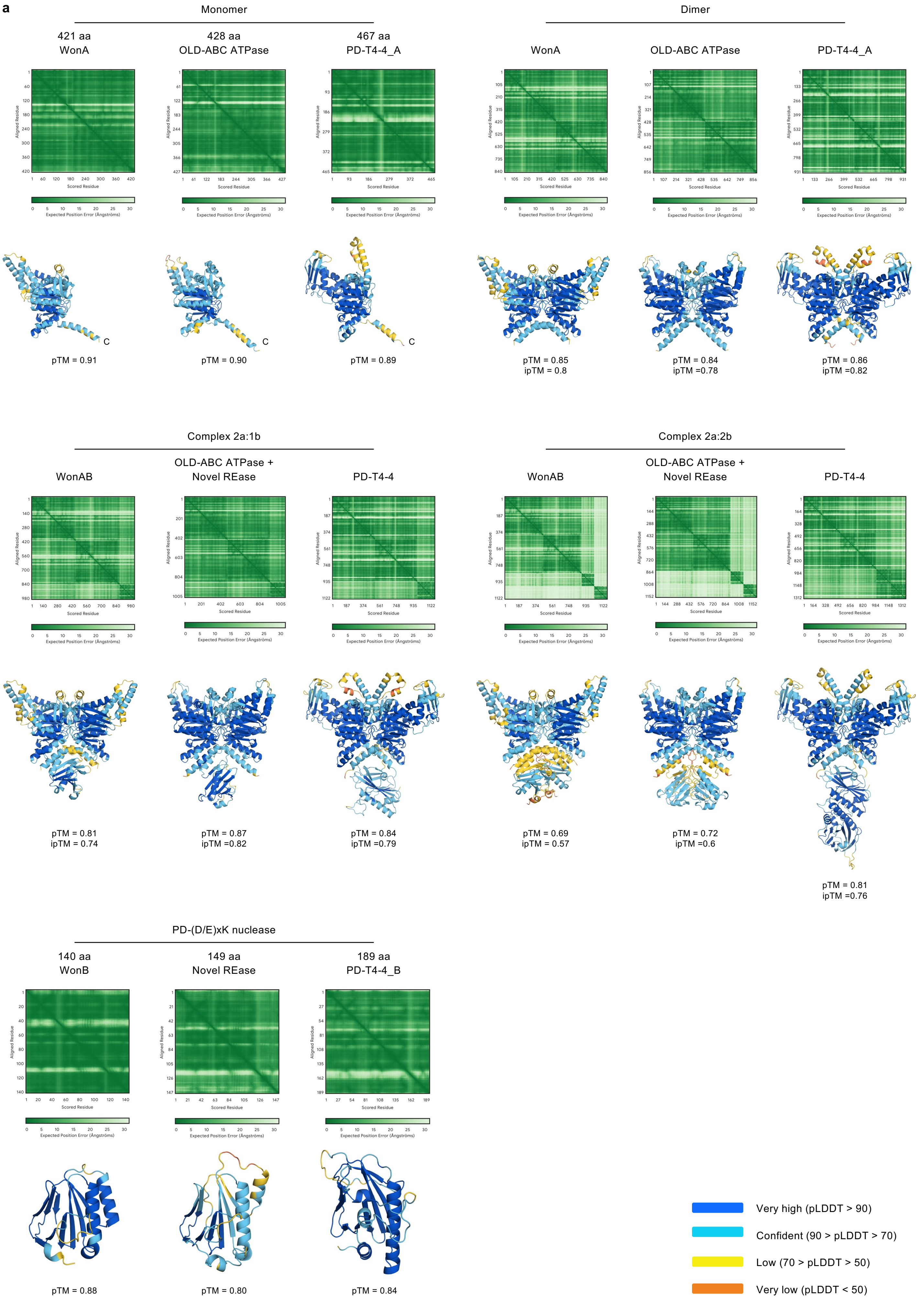

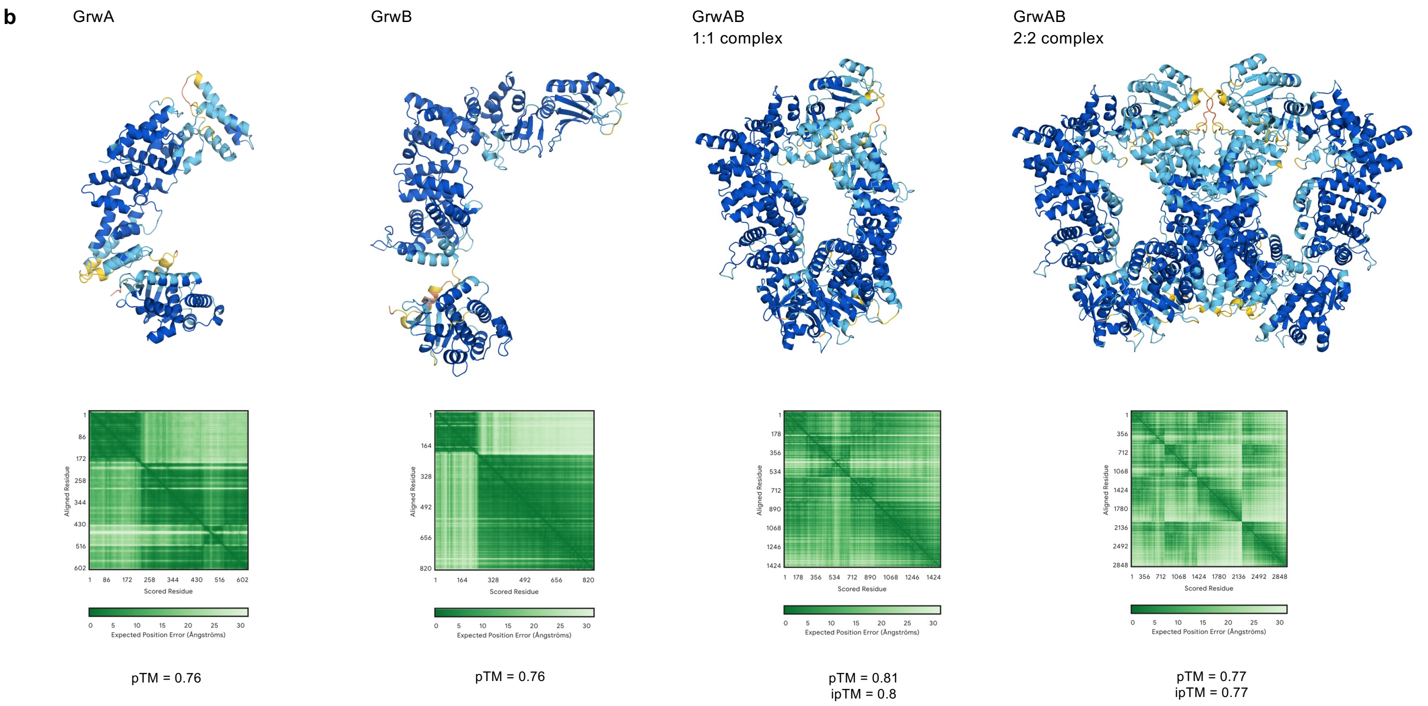

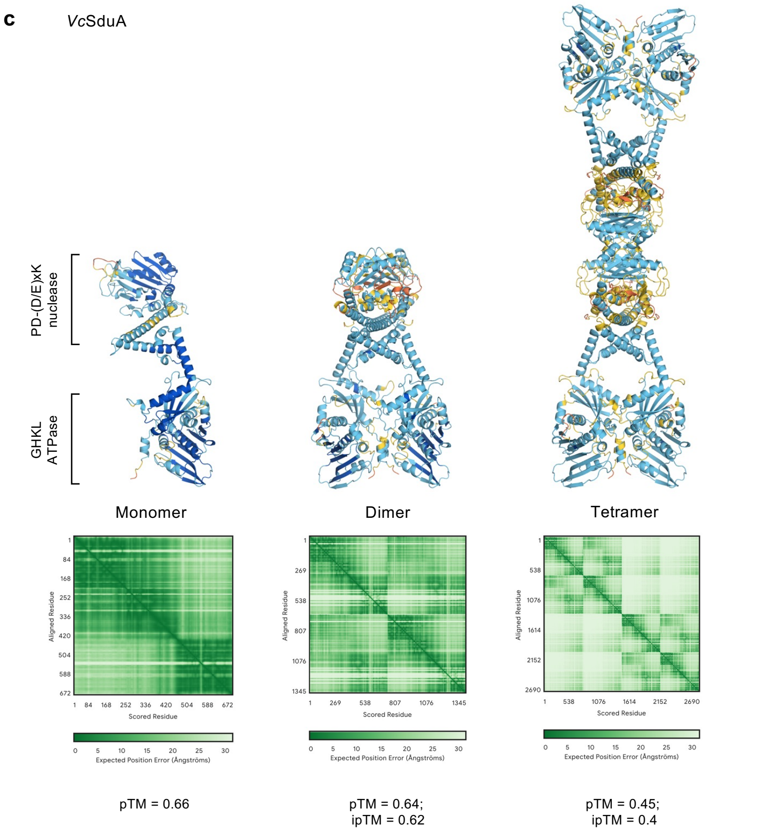

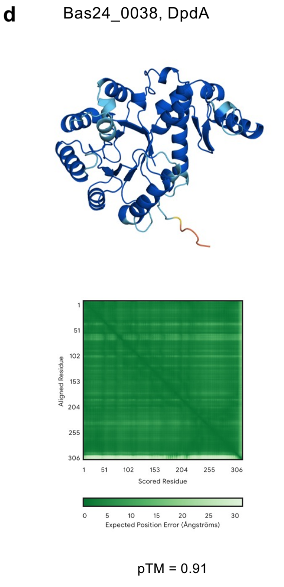
